# Viperin binds STING and enhances the type-I interferon response following dsDNA detection

**DOI:** 10.1101/493098

**Authors:** Keaton M. Crosse, Ebony A. Monson, Arti B. Dumbrepatil, Monique Smith, Yeu-Yang Tseng, Kylie H. Van der Hoek, Peter A. Revill, David C. Tscharke, E. Neil. G. Marsh, Michael R. Beard, Karla J. Helbig

**Affiliations:** Department of Physiology, Anatomy and Microbiology, La Trobe University, Bundoora, VIC, Australia; Departments of Chemistry and Biological Chemistry, University of Michigan, Ann Arbor, Ann Arbor, MI, United States; John Curtin School of Medical Research, The Australian National University, Canberra, ACT, Australia; School of Biological Sciences, The University of Adelaide, Adelaide, SA, Australia; Victorian Infectious Diseases Reference Laboratory, Royal Melbourne Hospital, Peter Doherty Institute for Infection and Immunity, Melbourne, VIC, Australia

**Keywords:** CIA2A, interferon, radical SAM enzyme, STING, viperin

## Abstract

Viperin is an interferon-inducible protein that is pivotal for eliciting an effective immune response against an array of diverse viral pathogens. Here we describe a mechanism of viperin’s broad antiviral activity by demonstrating the protein’s ability to synergistically enhance the innate immune dsDNA signalling pathway to limit viral infection. Viperin co-localised with the key signalling molecules of the innate immune dsDNA sensing pathway, STING and TBK1; binding directly to STING and inducing enhanced K63-linked polyubiquitination of TBK1. Subsequent analysis identified viperin’s necessity to bind the cytosolic iron-sulphur assembly component 2A, to prolong its enhancement of the type-I interferon response to aberrant dsDNA. Here we show that viperin facilitates the formation of a signalling enhanceosome, to coordinate efficient signal transduction following activation of the dsDNA signalling pathway; which results in an enhanced antiviral state. We also provide evidence for viperin’s radical SAM enzymatic activity to self-limit its immunomodulatory functions. This data further defines viperin’s role as a positive regulator of innate immune signalling, offering a mechanism of viperin’s broad antiviral capacity.

## Introduction

Innate immunity constitutes the first line of host defence against viral invasion, acting to both prevent and clear infection, as well as driving a robust adaptive response [1]. A range of germ-line encoded pattern recognition receptors (PRRs) are responsible for detecting various viral structural motifs, termed pathogen associated molecular patterns (PAMPs), upon which a signalling cascade is initiated to elicit an innate immune response [2, 3]. This response is primarily associated with the production of cytokines and chemokines, such as interferon (IFN), which acts in both an autocrine and paracrine manner to induce hundreds of interferon stimulated genes (ISGs) [4]. The products of these ISGs act to both clear the viral infection within the infected cell while simultaneously resisting infection in neighbouring cells (reviewed in [5]).

The induction of ISGs creates a highly effective antiviral state within a cell, yet the specific antiviral activity has only been characterised for a handful of these genes. Viperin (also known as cig5 and RSAD2) is one of the most well described and potent ISGs, implicated in limiting many viruses from multiple viral families (reviewed in [5, 6]). Viperin was first identified to have antiviral properties against human cytomegalovirus [5,7,8], and has since been shown to directly target multiple stages of the viral life cycle to inhibit infection of viruses including HCV, DENV, TBEV, HIV, BUNV, ZIKV and IAV (reviewed in [7, 9]). Viperin has been shown to target the budding of IAV, HIV-1 and RABV, by directly disrupting cellular lipid rafts through interactions likely involving the host protein FPPS [10–12]. Additionally, the replication of viruses such as HCV and DENV is also a target for viperin’s antiviral activity, whereby viperin has been demonstrated to associate with the viral non-structural proteins NS5A and NS3 respectively at the replication complexes of each virus to inhibit infection [13, 14]. However, there is not one direct means of viperin inhibition described for these multiple viral families, which all employ different routes of infection and mechanisms of replication (reviewed in [7, 15]). As such, viperin has been implicated in interactions with many functionally unrelated host and viral proteins, making it increasingly difficult to identify a unifying mechanism of viperin’s antiviral activity.

Viperin is a member of the radical SAM enzyme superfamily [16–19]. Recent work has demonstrated that mammalian viperin catalyses the dehydration of CTP to form 3’,4’-didehydro-4’-deoxy-CTP (ddhCTP) through a radical mechanism involving the reductive cleavage of SAM with concomitant formation of 5’-deoxyadenosine (5’-dA) [20]. Like all radical SAM enzymes, viperin relies for activity, on the insertion of a [4Fe-4S] cluster at the active site, which serves as the source of electrons in the reaction (Figure 1) [21, 22]. Cluster insertion is facilitated by the cytosolic iron-sulphur protein assembly (CIA) pathway, which comprises two distinct branches of targeting complexes [23]. Despite viperin’s interaction with both branches of CIA targeting complexes, only one has been shown to confer [4Fe-4S] insertion (Figure 1) [24].

**Figure 1.**
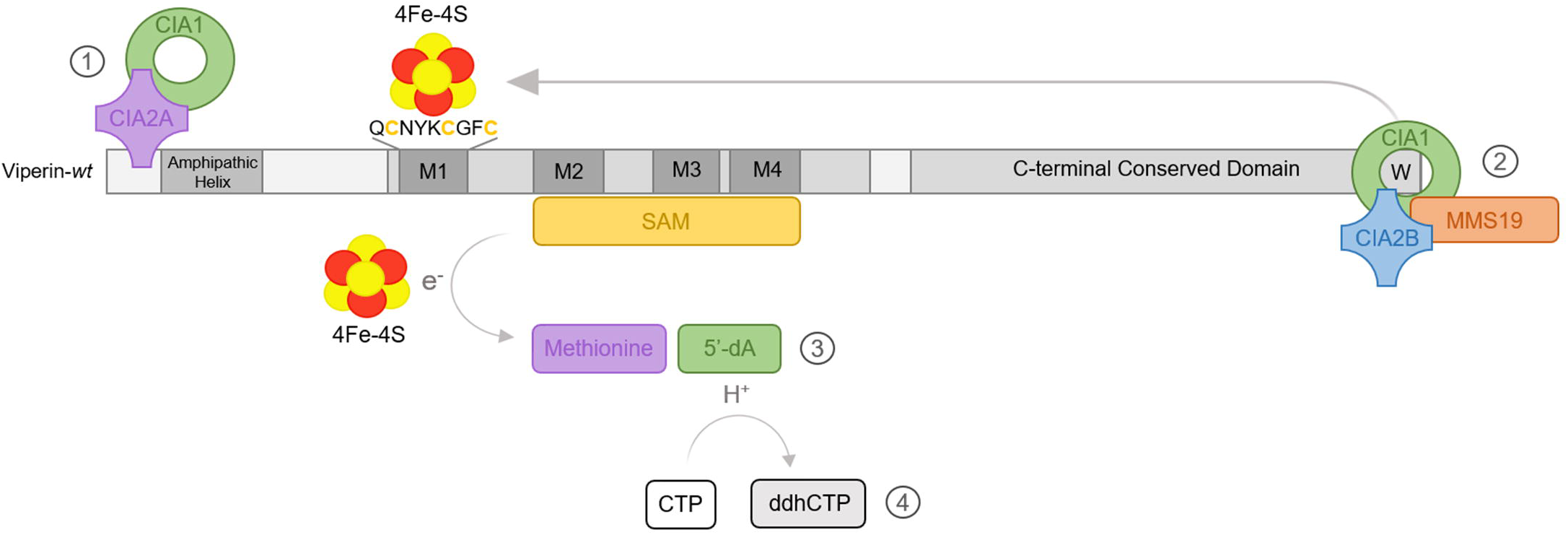
Schematic representation of viperin’s interaction with the CIA pathway, and enzymatic generation of ddhCTP. There are two distinct branches of the CIA pathway, each comprising a unique targeting complex, responsible for inserting Fe-S clusters into Fe-S apo-proteins. Each branch is categorised by the involvement of either isoform of the CIA2 protein, CIA2A or CIA2B, within the targeting complex. Generally, CIA2A is responsible for the stabilisation and maturation of proteins involved in iron homeostasis, while CIA2B enables the maturation of most cytosolic and nuclear Fe-S proteins involved in generic housekeeping [23]. 1) Viperin binds CIA2A-CIA1 complex within the first 50 amino acid residues of its N-terminus, but is not responsible for the insertion of [4Fe-4S] into viperin [23,24,45]. 2) The alternate CIA targeting complex CIA1-CIA2B-MMS19 binds to the conserved tryptophan residue (W361aa) at viperin’s C-terminus [45], and is responsible for [4Fe-4S] insertion into the conserved cysteine residues within viperin’s central M1 domain. 3) Viperin binds SAM at its central M2, M3 and M4 domains and catalyses its cleavage into methionine and 5’-Ado radical intermediate (5’-deoxyadeno<u>syl</u>), whereby an electron is abstracted by the bound [4Fe-4S] cluster. 4) The 5’-dAdo radical then abstracts a hydrogen from CTP, becoming 5’-dA (5’-deoxyadeno<u>sine</u>) and generating the antiviral ribonucleotide ddhCTP [20, 45].

ddhCTP has been shown to act as a chain terminator for viral RNA-dependent RNA polymerases (RdRp) from multiple *Flavivirus* members [20], and this offers an explanation for viperin’s ability to broadly limit this viral genus (reviewed in [7, 20]). However, ddhCTP is unable to inhibit the polymerase activities of the *Picornaviridae* members human rhinovirus (HRV) C and poliovirus, despite viperin’s previously identified antiviral capacity against HRV [25]; which highlights the possibility that this highly evolutionarily conserved antiviral host protein, may still be involved in other as yet unidentified antiviral mechanisms.

Viperin is one of a small group of ISGs that are capable of acting as both direct antiviral effectors and indirectly as enhancers of innate immune signalling, to inhibit viral infection (reviewed in [26]). Viperin can enhance the activation of key signalling molecules of both the TLR7 (ssRNA sensing) and TLR9 pathways (CpG DNA sensing), and is able to directly bind IRAK1 and TRAF6, to enhance the activation of IRAK1, via augmenting the molecule’s K63-linked polyubiquitination [27]. This in turn results in a heightened production of interferon, and an enhanced antiviral response. Multiple viruses have PAMPs that activate these pathways, including IAV and VSV [28, 29], which are known to be limited by viperin (reviewed in [7]), and this mechanism may help explain some of the broader antiviral activities of viperin.

Here we show for the first time that viperin is also able to directly interact with the signalling adaptor molecule, stimulator of IFN genes (STING), and enhance production of antiviral cytokines following activation of dsDNA signalling pathways, and that this relies not only on its radical SAM enzymatic activity, but also its interaction with the iron-sulfur assembly component 2A (CIA2A). Viperin’s ability to augment both dsDNA signalling, as well as TLR7 and 9 signalling, offers further explanation for this host protein’s ability to limit such a broad range of viral families.

## Results

### Viperin enhances the STING-dependent type-I IFN response to dsDNA downstream of ligand detection

To investigate whether viperin plays a role in the enhancement of the IFN response to dsDNA we initially utilised *in vitro* cell culture-based luciferase assays. Ectopic expression of viperin in both HeLa and Huh-7 cells was observed to enhance the activity of the type-I IFN-β promoter in dual luciferase reporter assays following stimulation of the DNA viral mimic, poly dA:dT by approximately 2.5-fold and 2-fold respectively (Figure 2Ai & ii). These results were confirmed with the use of our previously developed primary viperin^−/−^ MEFs [24] as well as a polyclonal Huh□7 cell line stably expressing shRNA targeting viperin mRNA [11]. As can be seen in Figure 2B, primary viperin^−/−^ MEFs displayed an approximate 4-fold reduction in their expression of IFN-β mRNA relative to wild-type MEFs, and the activity of the IFN-β promoter was significantly reduced in the shViperin cells compared to the shControl cell line following poly dA:dT stimulation (Figure 2C). Together this data demonstrates the ability of viperin to enhance the type-I IFN response to exogenous dsDNA stimulus in multiple cell lines.

**Figure 2.**
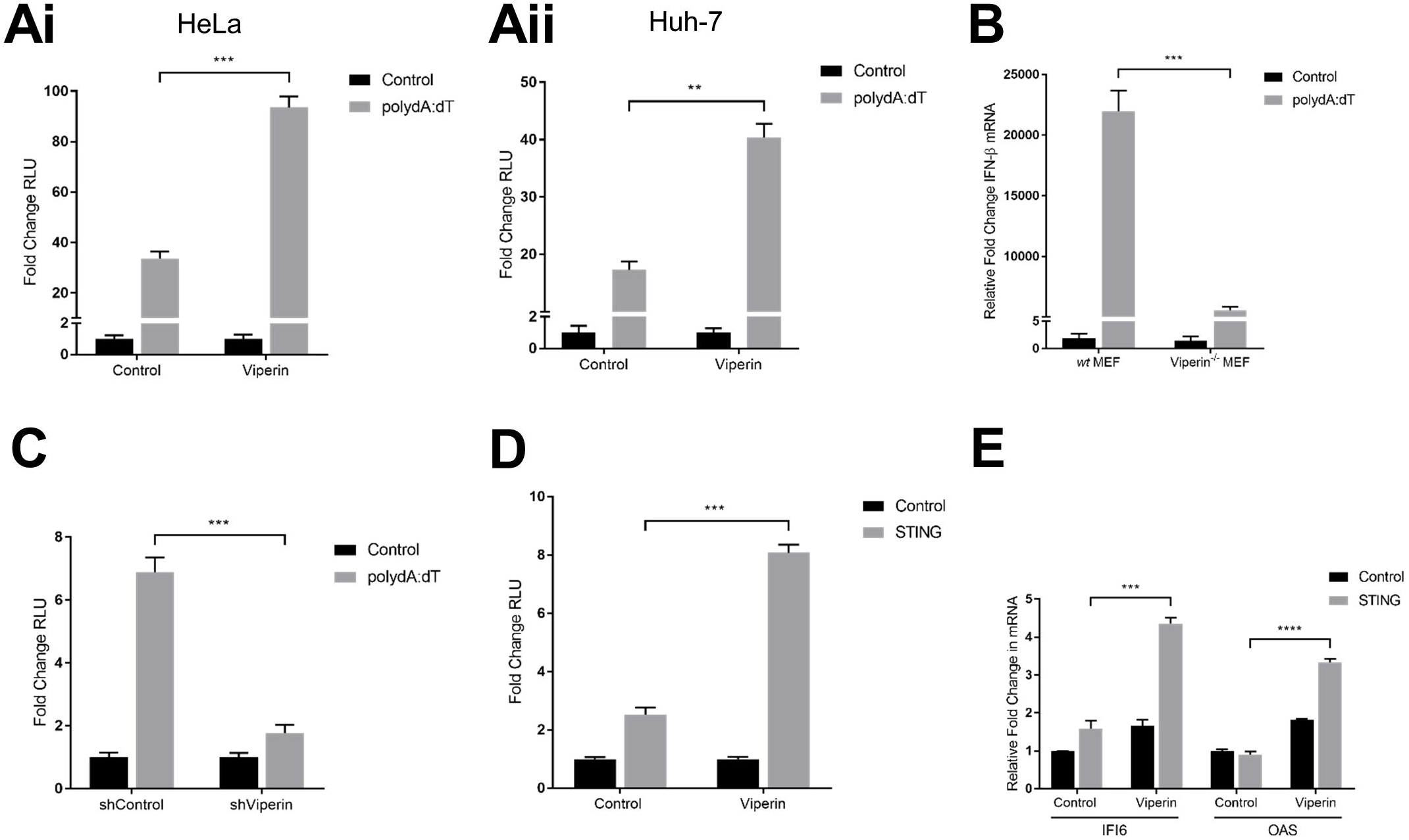
Viperin enhances STING-dependent type-I IFN response to dsDNA downstream of ligand detection. **(Ai & ii)** Luciferase production driven by the IFN-β promoter in HeLa **(i)** and Huh-7 **(ii)** cells transfected with either viperin or control constructs 24 hrs prior to stimulation with poly dA:dT for 8 hrs. **(B)** Expression of IFN-β mRNA in wild-type and viperin^−/−^ primary MEFs following 8 hr stimulation with poly dA:dT. **(C)** Luciferase production driven by the IFN-β promoter in Huh-7 cells stably expressing shRNA targeting viperin mRNA and its control stimulated with poly dA:dT for 8 hrs. **(D)** Luciferase production driven by the IFN-β promoter in Huh-7 cells transfected with combinations of viperin, STING or control constructs for 24 hrs. **(E)** Expression of IFI6 and OAS mRNA in HeLa cells transfected with combinations of viperin, STING or control constructs for 24 hrs. Luciferase measurements were controlled by constitutive expression of renilla and presented as fold changes in relative luminometer units (RLU) from control unstimulated conditions. All data is presented as mean ± SEM; *p<0.05, **p<0.01, ***p<0.001, ****p<0.0001; (n≥3).

The detection of exogenous dsDNA within the host cell relies on the activity of multiple DNA sensors, however upon recognition of their ligands these receptors predominantly converge on the adaptor molecule STING [30]. To identify whether viperin’s enhancement of the type-I IFN response to dsDNA stimulation involves an interaction with the downstream adaptor molecule STING, Huh-7 cells were co-transfected to ectopically express both viperin and STING in the absence of poly dA:dT stimulation. Cells expressing STING alone, in the absence of viperin expression, displayed a 3-fold increase in IFN-β promoter activity compared to cells transfected with control vector only (control)(Figure 2D), indicating that overexpression of this adaptor molecule is sufficient to auto-activate the pathway. Furthermore, in the absence of poly dA:dT stimulation, co-transfection of STING with viperin resulted in significantly higher IFN-β promoter activity compared to STING alone (Figure 2D), suggesting viperin’s enhancement of type-I IFN following dsDNA signalling occurs downstream of exogenous DNA recognition. Furthermore, viperin’s co-transfection with STING also significantly upregulated the production of the key antiviral interferon stimulated genes (ISGs) IFI6 and OAS compared to STING alone (Figure 2E), indicating that this positive augmentation of the type-I IFN pathway results in a functional upregulation of ISGs downstream of dsDNA ligand recognition.

### Viperin co-localises with TBK1 and STING, via a direct interaction with STING

The successful activation of signalling events initiated by dsDNA PRRs relies on the activity of two major adaptor molecules, STING and TBK1 [31, 32]. The ER-resident protein STING assembles with TBK1 after dsDNA stimulation to facilitate the phosphorylation of IRF3, culminating in the induction of type-I IFN [33, 34]. As viperin has previously been shown to co-localise and interact with alternate signalling adaptor molecules to enhance the efficacy of the TLR7/9 innate immune signalling pathways [27], we investigated viperin’s ability to co-localise with STING and TBK1.

Utilising *in situ* proximity localisation assays (PLA) in HeLa cells in conjunction with ectopically expressed viperin, we observed the co-localisation of viperin with endogenous STING and to a lesser degree endogenous TBK1 irrespective of poly dA:dT stimulation (Figure 3A). This co-localisation observed between viperin and TBK1/STING appeared to be enhanced during poly dA:dT stimulation (Figure 3A). To confirm these observations, we utilised confocal microscopy, where similar co-localisation was observed between viperin and either TBK1 or STING, however considerable co-localisation was only observed between viperin and TBK1 or STING following dsDNA stimulation (Figure 3B and C). As can be seen in Figure 3B, the cytoplasmic localisation of TBK1 appeared to converge with viperin on lipid droplets, 2 hrs following poly dA:dT stimulation. Similar to viperin’s localisation with TBK1, viperin appears to co-localise with STING around the BODIPY-stained lipid droplets following poly dA:dT stimulation (Figure 3C, indicated by white arrow), however at this point there is also considerable co-localisation of viperin and STING at discrete puncta throughout the cytoplasm, which is a hallmark of STING activation on the golgi [35].

**Figure 3.**
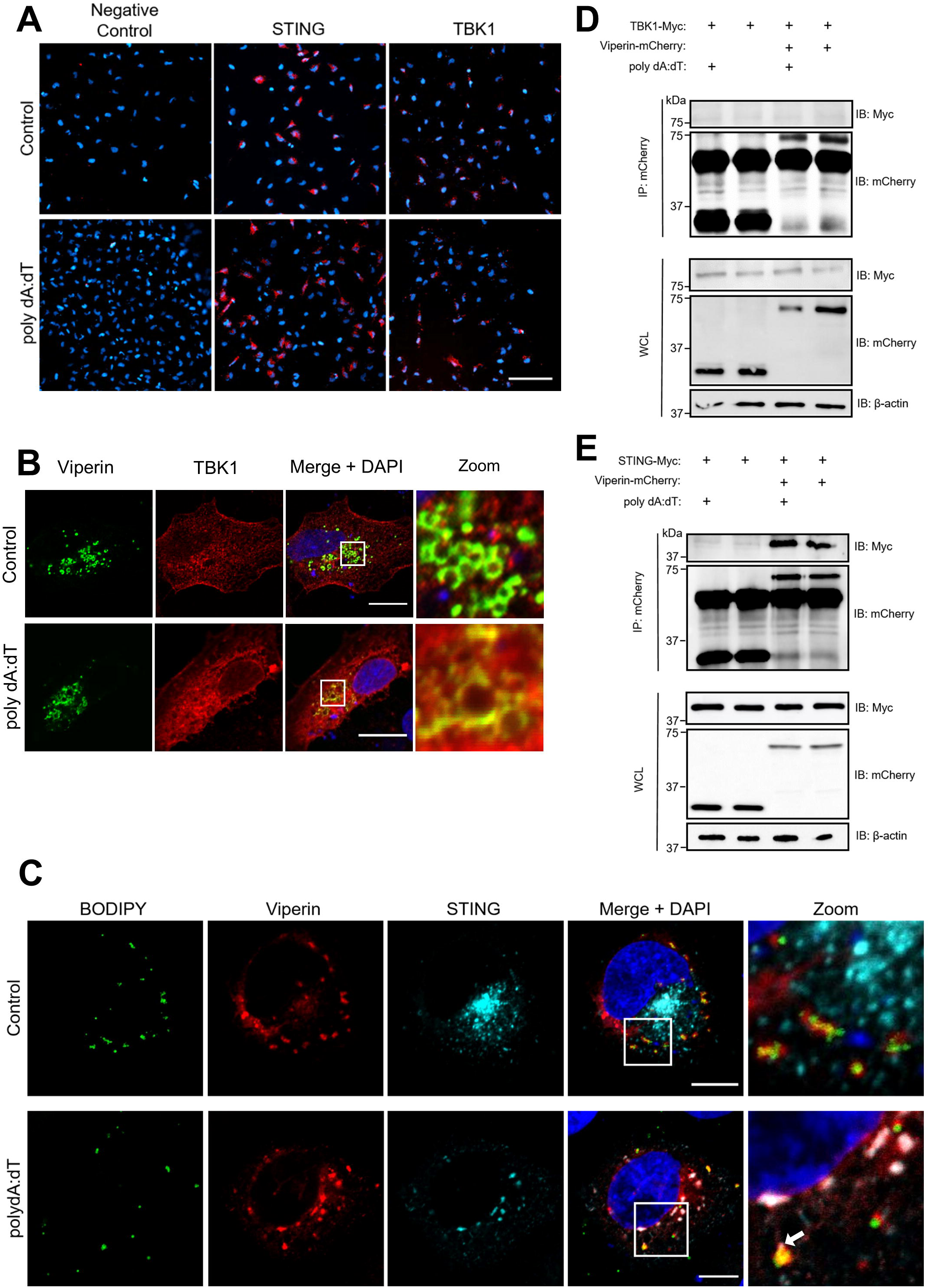
Viperin co-localises with TBK1 and STING, via a direct interaction with STING. **(A)** HeLa cells were transfected with a viperin-flag construct 24 hrs prior to stimulation with poly dA:dT for 2 hrs and probing with mouse monoclonal anti-flag (Sigma) and rabbit monoclonal anti-STING (Millipore) or anti-TBK1 (Cell Signalling) antibodies, then subject to Duolink® *In Situ* Red Mouse/Rabbit PLA and DAPI staining. Imaged on Nikon Eclipse T*i*-E fluorescence inverted microscope. Scale bar represents 150 µm. Original magnification is X20. **(B)** Huh-7 or **(C)** HeLa cells were transfected with viperin-flag and either **(B)** TBK1-myc or **(C)** STING-myc constructs 24 hrs prior to stimulation with poly dA:dT for 2 hrs and immunofluorescence staining with rabbit monoclonal anti-flag (Sigma) and mouse monoclonal anti-myc (Millipore) antibodies followed by an Alexa555-conjugated goat anti-mouse (Invitrogen) and Alexa488-conjugated goat anti-rabbit (Invitrogen) secondary, as well as DAPI and **(C)** BODIPY staining. Imaged on Ziess Confocal LSM 780 microscope. Scale bar represents 15 µm. Original magnification is X63. **(D and E)** Huh-7 cells were transfected with viperin-mCherry or control-mCherry and either **(D)** TBK1-myc or **(E)** STING-myc constructs 24 hrs prior to stimulation with poly dA:dT for 2 hrs, and cell extracts were immunoprecipitated with rabbit monoclonal anti-mCherry antibody (Biovision) and subject to immunoblot analysis with indicated antibodies. **(D)** Cell extract was subject to DSS crosslinking prior to lysis and immunoprecipitation. Immunoblots are representative of at least 2 independent experiments.

To further investigate the ability of viperin to form a complex with either STING or TBK1, co-immunopreciptation assays were performed. Preliminary immunoblot analysis was unable to detect TBK1 with immunoprecipitated viperin (Supplementary Figure 1A). To elucidate whether this was the result of potentially weaker or transient binding interactions between the two proteins, a DSS cross-linker was utilised. Despite the addition of the cross-linker, TBK1 failed to be co-immunoprecipitated with viperin (Figure 3D). However, STING was successfully detected following co-immunoprecipitation assays with viperin, irrespective of poly dA:dT stimulation (Figure 3E). Further analysis identified the central domain of viperin to be responsible for this binding to STING (Supplementary Figure 1B). Together these findings highlight the strong interaction between viperin and STING, and imply the formation of a complex between these two proteins, while also indicating that the co-localisation between viperin and TBK1 does not involve direct binding.

### Viperin enhances the polyubiquitination-dependent activation of TBK1

The adaptor molecules involved in innate immune signalling events are commonly regulated by post-translational modifications such as polyubiquitination [36]. The addition of both K27- and K63-linked ubiquitin chains to STING has been shown to facilitate optimal trafficking of the protein, enabling efficient activation of downstream signalling adaptors [37, 38]. To delineate viperin’s mechanism of enhanced IFN-β promoter activity in the presence of dsDNA, we first investigated STING activation in the presence of viperin. However, viperin expression was not found to impact either K27- or K63-linked polyubiquitination of STING in HEK293T cells (Figure 4A and B). Moreover, the presence of dimerised and phosphorylated forms of STING, which are associated with the protein’s ligand binding affinity and recruitment of downstream adaptor molecules respectively [39, 40], were unaffected by the co-expression of viperin in HEK293T cells (Figure 4C).

**Figure 4.**
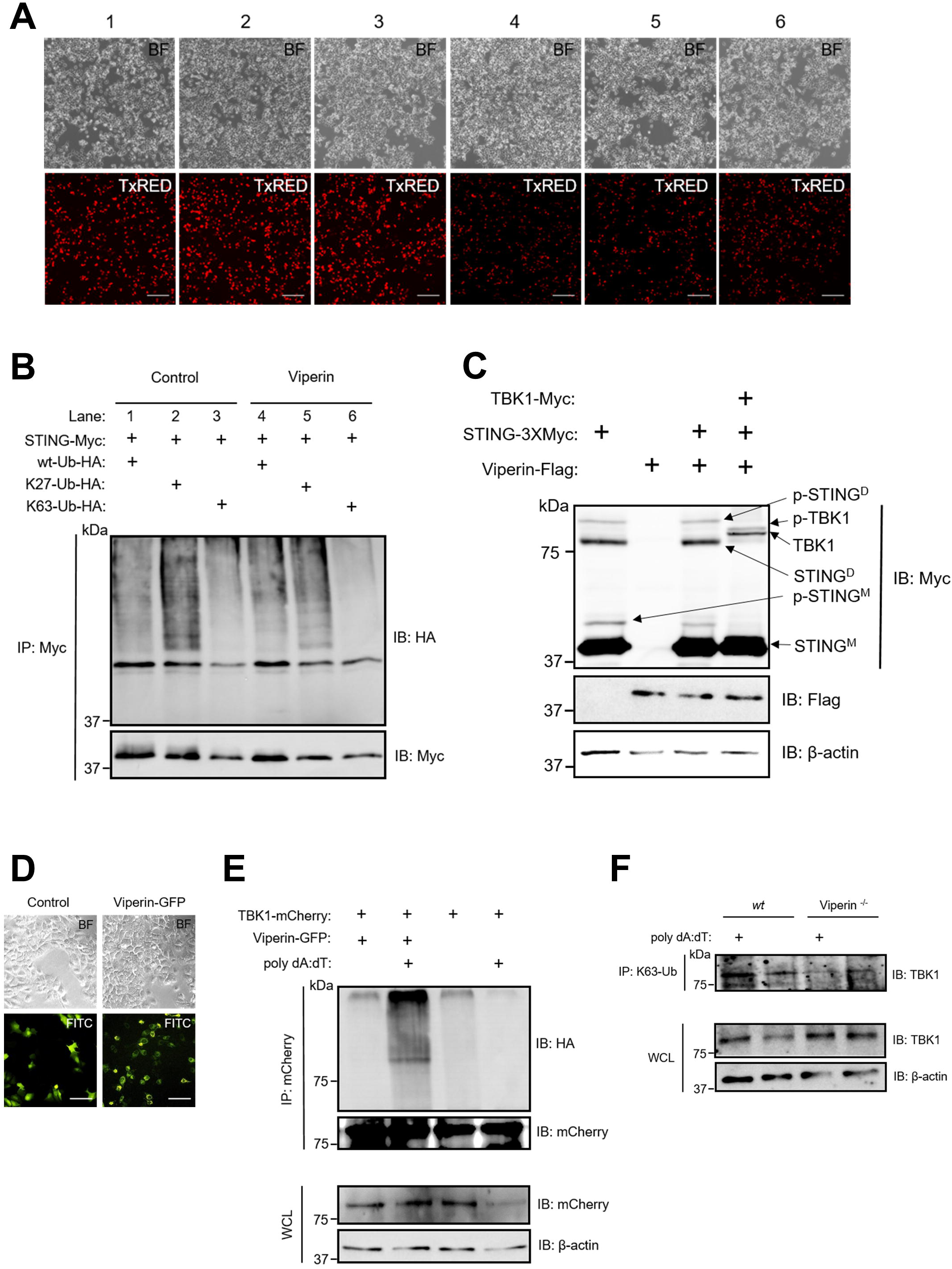
Viperin enhances the polyubiquitination-dependent activation of TBK1. **(A and B)** HEK293T cells were transfected with viperin-mCherry, STING-myc and either K63-Ub-HA, K27-Ub-HA or wt-Ub-HA constructs 24 hrs prior to **(A)** visualisation by fluorescence microscopy, after which **(B)** cell extracts were immunoprecipitated with mouse monoclonal anti-myc antibody (Millipore) and subject to immunoblot analysis with indicated antibodies. Imaged on Nikon Eclipse T*i*-E fluorescence inverted microscope. Scale bar represents 200 µm. Original magnification is X20. **(C)** HEK293T cells were transfected with combinations of viperin-flag, STING-myc and TBK1-myc constructs 24 hrs prior to immunoblot analysis with indicated antibodies. **(D and E)** Huh-7 cells were transfected with viperin-GFP, TBK1-mCherry and K63-Ub-HA constructs 24 hrs prior to **(D)** visualisation by fluorescence microscopy, and **(E)** stimulation with poly dA:dT for 2 hrs, after which cell extracts were immunoprecipitated with rabbit monoclonal anti-mCherry antibody (Biovision) and subject to immunoblot analysis with indicated antibodies. Imaged on Nikon Eclipse T*i*-E fluorescence inverted microscope. Scale bar represents 100 µm. Original magnification is X20. **(F)** Wild-type and viperin^−/−^ primary MEFs were stimulated with poly dA:dT for 2 hrs, and cell extracts were immunoprecipitated with mouse monoclonal anti-K63-Ub antibody (Enzo) and subject to immunoblot with indicated antibodies. Immunoblots are representative of at least 2 independent experiments.

The activation of TBK1 is regulated through the conjugation of K63-linked ubiquitin chains to lysine residues 30 and 401 along the protein [41]. To determine the impact of viperin on this polyubiquitination event, ectopically expressed viperin was visualised in Huh-7 cells (Figure 4D), prior to the determination of the polyubiquitination status of TBK1 through immunoprecipitation coupled with immunoblot analysis. There was substantial K63-linked polyubiquitination of TBK1 observed in cells containing viperin following a 2 hr poly dA:dT stimulation (Figure 4E), in contrast to the control. Furthermore, in primary wild-type MEFs, K63-linked polyubiquitination of endogenous TBK1 was observed to be markedly increased following poly dA:dT stimulation, compared to viperin^−/−^ MEFs (Figure 4F). Collectively this data demonstrates that viperin enhances the K63-linked polyubiquitination of TBK1.

### Viperin interacts with STING to enhance the type-I IFN response to limit HBV

To investigate whether viperin’s ability to enhance a type-I IFN response to dsDNA would functionally affect the outcome of a DNA viral infection, we utilised a well-characterised HBV *in vitro* model viral system [42]. A 1.3 mer HBV plasmid transfection model for two prevalent HBV genotypes (HBV-D & HBV-A) was utilised in HepG2 cells ectopically expressing a combination of viperin and STING.

To determine the involvement of viperin in eliciting a type-I IFN response, the above-mentioned HBV infection model was utilised in conjunction with a dual luciferase reporter assay. At 48 hrs post transfection with either HBV-D or HBV-A, but not the control plasmid, cells expressing both viperin and STING showed an approximate 20-fold and 60-fold increase respectively in the induction of IFN-β compared to those only expressing STING (Figure 5A); suggesting that the interaction between viperin and STING drives an enhanced type-I interferon response to HBV infection.

**Figure 5.**
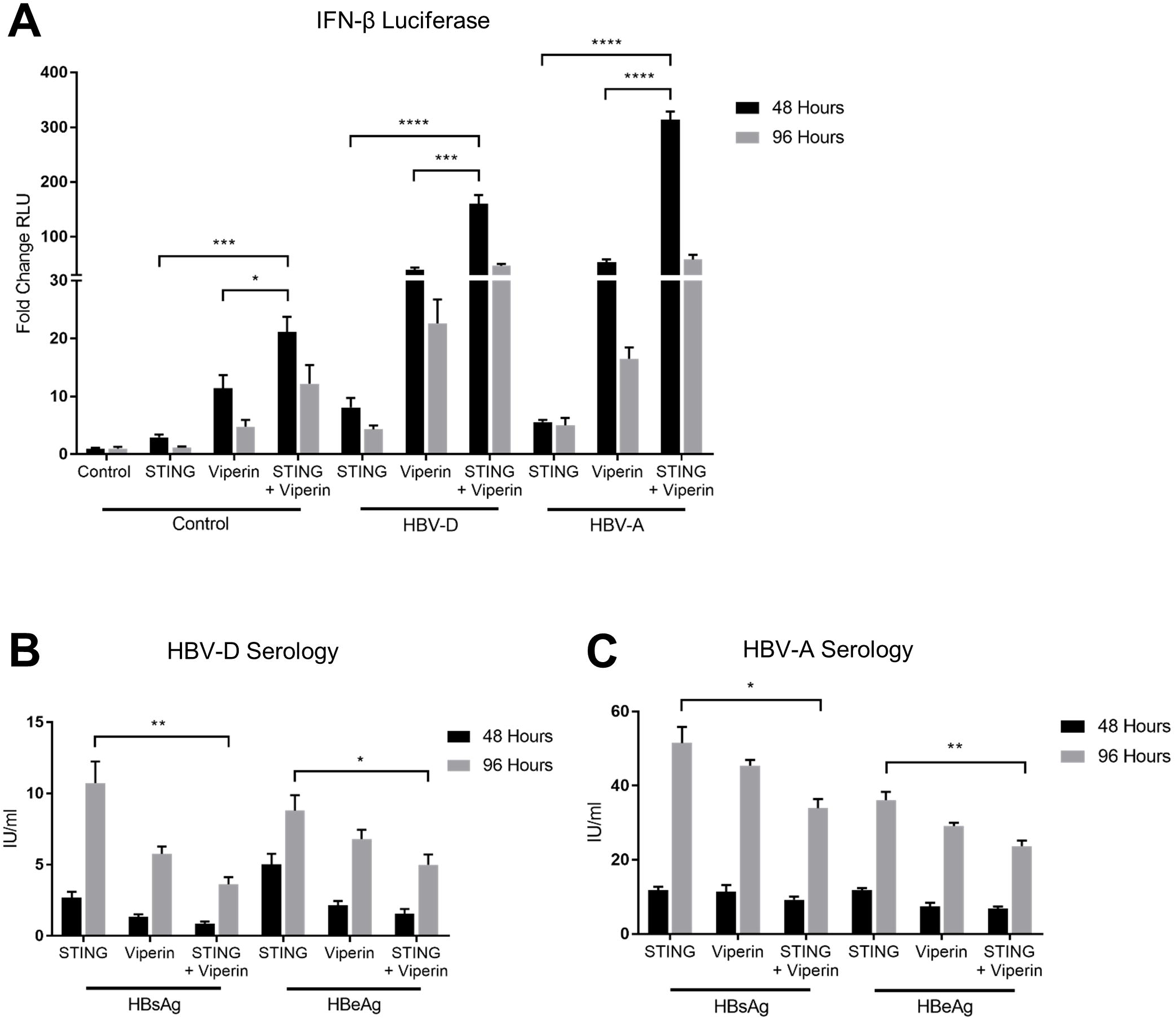
Viperin interacts with STING to enhance the type-I IFN response to limit HBV. **(A, B and C)** HepG2 cells were transfected ectopically with viperin, STING and/or control plasmids in conjunction with either HBV-D or HBV-A 1.3 mer constructs 48 or 96 hrs prior to lysis and; **(A)** IFN-β promoter driven luciferase production was detected, while **(B and C)** supernatants were collected for detection of HBsAg and HBeAg by Roche Cobas Elecsys quantitative serology for both HBV-D and HBV-A genotypes (n=4). Data is presented as mean ± SEM; *p<0.05, **p<0.01, ***p<0.001, ****p<0.0001; (n≥3).

To evaluate the effect of the enhanced type-I IFN response to HBV infection in the presence of viperin, cell supernatants were collected 48 and 96 hrs post HBV transfection and analysed by quantitative serology for the presence of HBV surface antigen (HBsAg) and HBV e antigen (HBeAg). At 96 hrs post HBV-A transfection, the ectopic expression of viperin in combination with STING significantly reduced the presence of both HBsAg and HBeAg circulating in cell supernatants, compared to cells either expressing either STING or viperin alone (Figure 5B). Similarly, a significant reduction in HBsAg and HBeAg was also observed in supernatants derived from cells expressing both viperin and STING compared to those solely expressing STING or viperin when transfected with the genotype-D HBV for 96 hrs (Figure 5C). STING activation has previously been demonstrated to limit HBV replication [43, 44], and collectively, these results demonstrate that a STING mediated innate response can be enhanced by viperin, to control HBV infection *in vitro*.

### Viperin’s N-terminus and [[4Fe-4S]] cofactor contribute to its enhancement of the type-I IFN response to dsDNA

Viperin relies on the action of specific functional domains to inhibit multiple families of viral pathogens [7]. Its characteristic localisation to the outer lipid droplet membrane (Figure 6A) is essential for its ability to enhance the type-I IFN response via TLR7/9 activation of plasmacytoid dendritic cells in the mouse [27]. To determine the potential role of certain viperin domains in its ability to enhance type-I IFN following poly dA:dT stimulation we utilised *in vitro* luciferase assays in combination with a panel of viperin mutants. As can be seen in Figure 6B, the 5’Δ33 viperin mutant, which lacks the first 33 amino acids of the N-terminus, and hence loss of its amphipathic helix [14], and the SAM1 viperin mutant in which the [4Fe-4S] cluster-binding cysteine residues of the protein’s radical SAM Motif 1 are mutated to alanine [14], both showed a significant decrease in IFN-β promoter activity compared to those expressing viperin-wildtype following poly dA:dT stimulation, and resembled that of the cells entirely lacking viperin (control). Conversely, the 3’Δ17 viperin mutant which lacks 17 amino acids from the protein’s C-terminus [14], significantly enhances viperin’s ability to upregulate IFN-β promoter activity (Figure 6B). This would imply that viperin requires either its localisation to the lipid droplet or specific sequences within its N-terminus, in conjunction with its binding to the [4Fe-4S] cluster, to enhance the type-I IFN response to dsDNA. Additionally, the ability of the 3’Δ17 viperin mutant to maintain enhancement of the IFN-β promoter driven luciferase, indicates that viperin does not require its radical SAM activity to augment the interferon response following detection of dsDNA, as the truncated C-terminal amino acids are required for [4Fe-4S] loading, a requirement for its enzymatic function [45].

**Figure 6.**
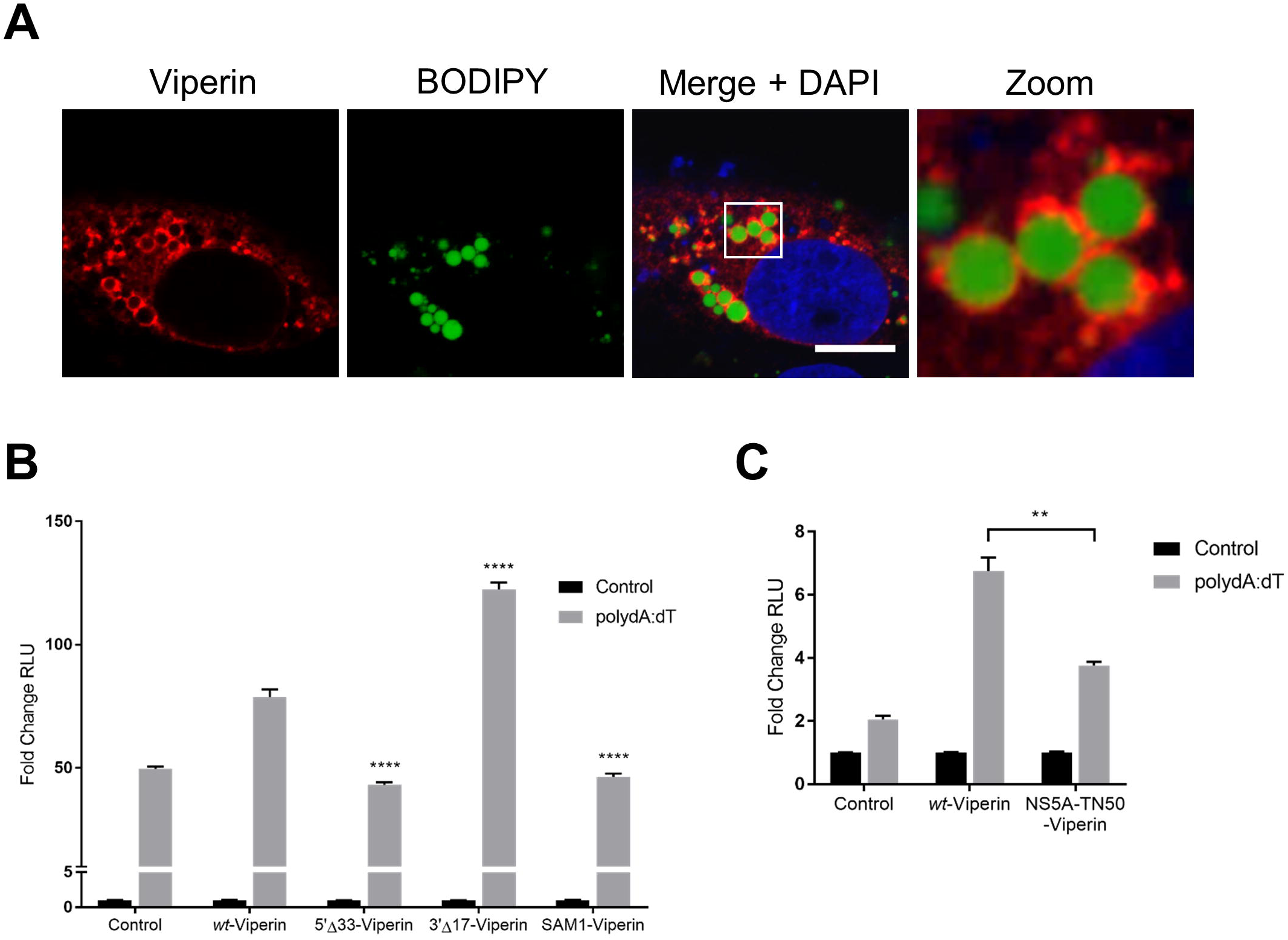
Viperin relies on its N-terminus and binding to enzymatic [4Fe-4S] cofactor to enhance type-I IFN response to dsDNA. **(A)** Huh-7 cells were transfected with viperin-flag 24 hrs prior to immunofluorescence staining with a mouse monoclonal anti-flag antibody (Sigma), followed by an Alexa555-conjugated goat anti-mouse secondary (Invitrogen) as well as BODIPY and DAPI staining. Imaged on Ziess Confocal LSM 780 microscope. Scale bar represents 10 µm. Original magnification is X63. **(B and C)** Luciferase production driven by the IFN-β promoter in HeLa cells transfected with either **(B)** wild-type, 5’Δ33, 3’Δ17 and SAM1 viperin constructs or **(C)** wild-type and chimeric NS5A-viperin mutant constructs 24 hrs prior to stimulation with poly dA:dT for 8 hrs. Luciferase measurements were controlled by constitutive expression of renilla and presented as fold changes in relative luminometer units (RLU) from control unstimulated conditions. Data is presented as mean ± SEM; **p<0.01, ****p<0.0001; (n≥3).

To further investigate the requirement of viperin’s N-terminus in its augmentation of the dsDNA signalling pathway, we utilised a chimeric viperin mutant (NS5A-TN50-Viperin). This mutant lacks 50 amino acids from viperin’s N-terminus, including its lipid droplet-localising amphipathic helix, but is substituted with the alternate amphipathic helix of HCV NS5A [45, 46]. Despite retaining viperin’s typical cellular localisation [45], this mutant was unable to enhance the induction of the IFN-β promoter to the same degree as wild-type viperin (p<0.01) (Figure 6C) following poly dA:dT stimulation, suggesting that localization to the lipid droplets is not of itself, sufficient for viperin’s enhancement of the type-I IFN response to dsDNA and there may be additional N-terminal sequences contributing to this activity.

### Viperin’s interaction with STING and TBK1 activates it towards the synthesis of ddhCTP and facilitates self-limiting degradation

As a radical SAM enzyme, viperin couples the reductive cleavage of SAM to the dehydration of CTP producing 5’-dA and ddhCTP as products (Figure 1) [20]. We have previously shown that viperin’s interaction with IRAK1 and TRAF6 to enhance TLR7/9 signalling results in significant stimulation of the enzyme’s catalytic activity. Interestingly, the interaction with IRAK1 and TRAF6 resulted in more rapid degradation of viperin, which would act to limit viperin’s antiviral activity [20, 47]. Moreover, high levels of 5’-dA have been demonstrated to act as a general inhibitor of radical SAM enzymes [48]. Therefore, we assessed whether viperin’s interaction with the signalling adaptor molecules STING and TBK1 would similarly stimulate viperin’s catalytic activity and result in more rapid degradation of viperin. The enzymatic activity of viperin was assayed in extracts prepared from HEK293T cells expressing either viperin and/or STING and TBK1, as described previously [47]. Consistently, the overexpression of STING, and to a lesser degree TBK1, significantly increased the catalytic activity of viperin, as determined by the amount of 5’-dA formed in the assay, compared to viperin alone when normalized for viperin expression levels (Figure 7A). Interestingly, through immunoblot analysis we also observed significant degradation of viperin in HEK293T cells overexpressing STING at 40 hrs post transfection, which was not observed at earlier time points (Figure 7B). Together these data demonstrate that viperin’s interaction with the signalling adaptor molecules of the dsDNA signalling pathway, STING and TBK1, both enhance viperin’s enzymatic activity and facilitate viperin’s degradation. STING and TBK1 thereby act as a negative feedback loop to limit viperin’s enhancement of immune signalling.

**Figure 7.**
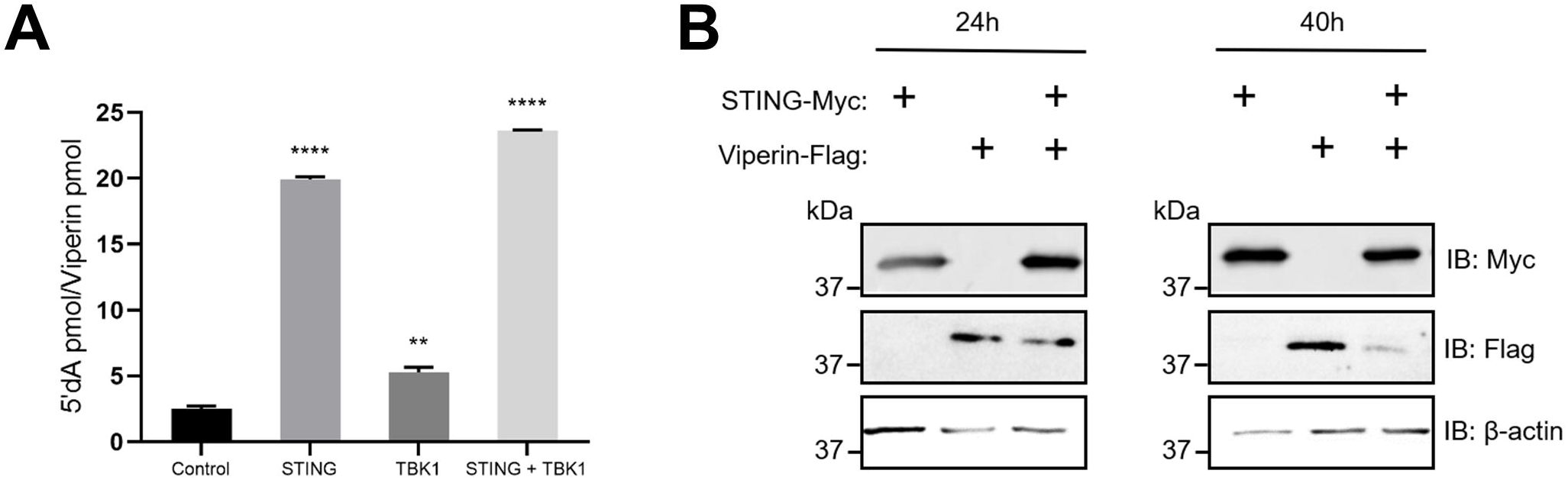
Viperin’s interaction with STING and TBK1 drives its radical SAM enzymatic activity and facilitates self-limiting degradation. **(A)** Quantification of viperin activity in cell extracts. HEK293T cells were transfected with combinations of viperin-flag as well as STING-myc and/or TBK1-myc constructs prior UPLC-tandem mass spectrometry quantification of 5’-dA generated in 1 h following addition of CTP and SAM. 5’-dA levels are presented relative to levels of viperin, as determined by quantitative immunoblot analysis. Data is presented as mean ± SEM; **p<0.01, ****p<0.0001; (n=6). **(B)** HEK293T cells were transfected with combinations of viperin-flag and STING-myc constructs 24 and 40 hrs prior to immunoblot analysis with indicated antibodies (The 24 hr immunoblot is a portion of figure 4C for reference). Immunoblots are representative of at least 2 independent experiments.

### Viperin co-localises with CIA2A to enhance the type-I IFN response to dsDNA

The N-terminal region of viperin is responsible for its binding to the cytosolic iron-sulphur protein assembly (CIA) component 2A (CIA2A) (Figure 1), a protein which is part of a pathway responsible for the targeting of Fe-S clusters to some proteins, but has been reported to not contribute to the insertion of the [[4Fe-4S]] cluster into viperin [24]. Therefore, given viperin’s requirement for N-terminal sequences to positively augment the type-I IFN response to dsDNA detection, we investigated a potential role for CIA2A in this novel viperin function.

Confocal microscopy revealed a cytoplasmic and nucleoplasmic localisation of CIA2A, which did not show any notable co-localisation with BODIPY-stained lipid droplets (Figure 8A). However, the expression of viperin redistributed the localisation of CIA2A to convene on the lipid droplets, where the two proteins co-localised (Figure 8A). To determine the role of CIA2A in viperin’s enhancement of the IFN response to dsDNA we utilised CRISPR/Cas9 technology to generate CIA2A-deficiencies in HeLa cells. We verified loss of CIA2A in three independent polyclonal populations of HeLa cells, each containing a distinct gRNA (hereafter referred to as CIA2A KO #1, CIA2A KO #2 and CIA2A KO #3)(Figure 8B). Interestingly, these cells all displayed defects in their ability to drive an IFN-β promoter generated lucferase response when challenged with polydA:dT (Figure 8C). Most notably, the viperin-mediated enhancement of the IFN-β promoter activity was significantly reduced in all three of the CIA2A-deficient cell populations compared to wild-type HeLa cells (Figure 8C). Conversely, the combined overexpression of CIA2A and viperin further enhanced the activity of the IFN-β promoter following polydA:dT stimulation by 3-fold compared to viperin alone (Figure 8D). These findings implicate viperin’s N-terminal association with CIA2A as a significant contribution towards its ability to enhance the type-I IFN response to dsDNA.

**Figure 8.**
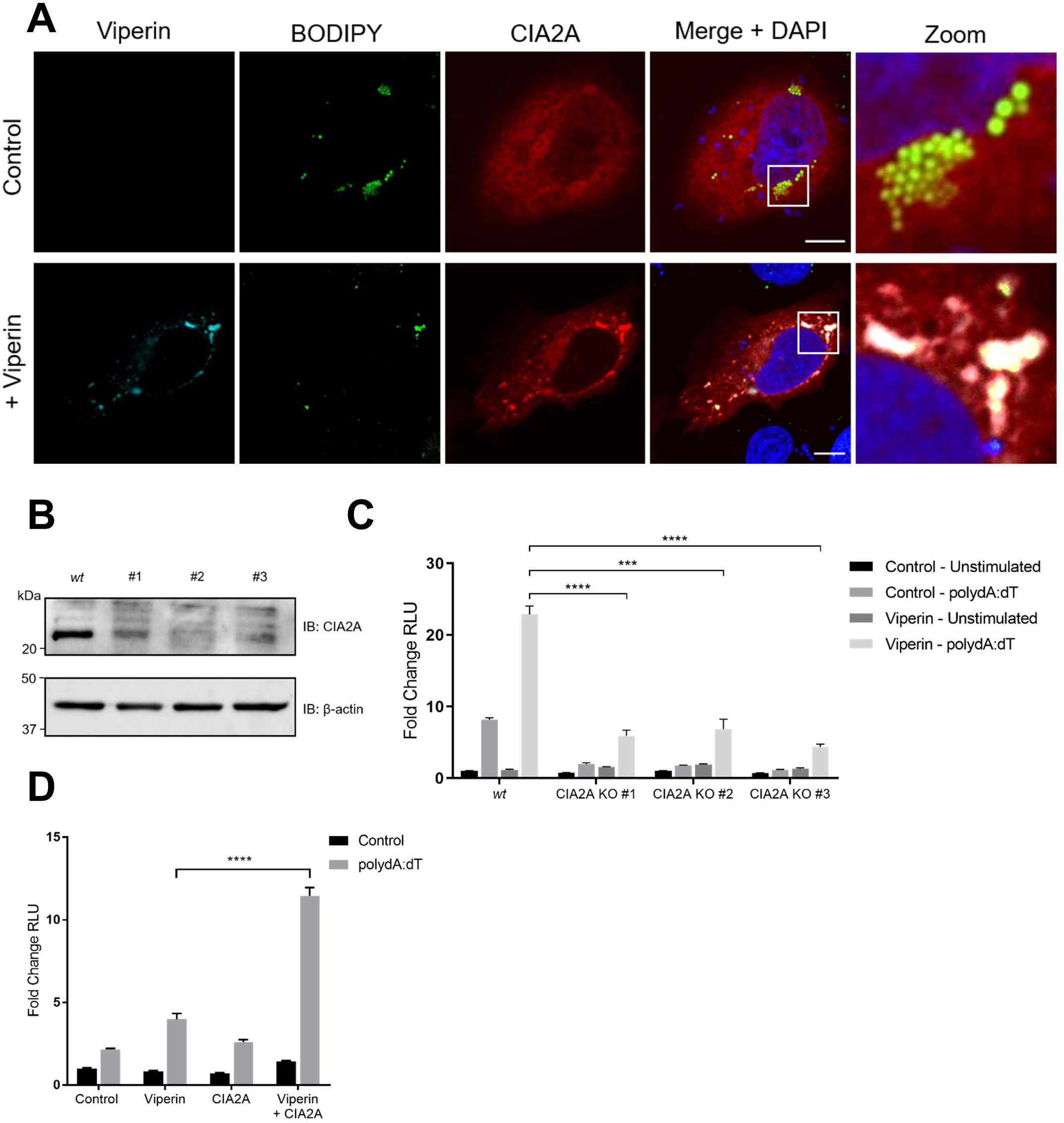
Viperin co-localises with CIA2A to enhance the type-I IFN response to dsDNA. **(A)** HeLa cells were transfected with viperin-flag and/or CIA2A-myc constructs 24 hrs prior to immunofluorescence staining with a rabbit monoclonal anti-flag antibody (Sigma) and/or mouse monoclonal anti-myc (Millipore), followed by an Alexa555-conjugated goat anti-rabbit (Invitrogen) and/or Alexa647-conjugated goat anti-mouse (Invitrogen) secondaries as well as BODIPY and DAPI staining. Imaged on Ziess Confocal LSM 780 microscope. Scale bar represents 10 µm. Original magnification is X63. **(B)** Immunoblot analysis of CIA2A expression in HeLa wild-type (Cas9) and CRISPR-Cas9 polyclonal CIA2A knockdown cells; immunoblots are representative of 2 independent experiments. **(C)** Luciferase production driven by the IFN-β promoter in HeLa wild-type (Cas9) and CRISPR-Cas9 polyclonal CIA2A knockdown cells transfected with viperin constructs 24 hrs prior to stimulation with poly dA:dT for 8 hrs. **(C)** Luciferase production driven by the IFN-β promoter in HeLa cells transfected with viperin and/or CIA2A constructs 24 hrs prior to stimulation with poly dA:dT for 8 hrs. Luciferase measurements were controlled by constitutive expression of renilla and presented as fold changes in relative luminometer units (RLU) from control unstimulated conditions. Data is presented as mean ± SEM; ***p<0.001, ****p<0.0001; (n≥3).

## Discussion

Viperin is a potent antiviral host protein, associated with the inhibition of a broad range of viral infections (reviewed in [7]), however the scope of viperin’s antiviral capacity makes it increasingly difficult to discern the protein’s mechanism of viral inhibition. To date viperin has been shown to target multiple stages of viral lifecycles through interactions with many, often functionally unrelated, host and viral proteins (reviewed in [7]). The recent discovery that viperin synthesizes the novel nucleotide, ddhCTP, offers an explanation for its ability to inhibit *Flavivirus* infection and unequivocally links the protein’s enzymatic function in its antiviral activity [20]. However, this mechanism of viperin’s antiviral activity fails to account for each instance of the protein’s ability to limit diverse viral pathogens. Recent literature has suggested that viperin’s ability to positively regulate innate immune responses may elucidate a unifying antiviral mechanism for this potent antiviral protein [26, 27]. Here we show for the first time that viperin is able to enhance the innate immune response to dsDNA viral mimics and to DNA viral infection independently of its radical SAM activity, but dependent on the cytosolic iron-sulfur assembly component 2A (CIA2A). This work adds to the limited knowledge of viperin’s alternate innate immune regulatory capacity.

Viperin’s enhancement of the cellular innate response to dsDNA shares mechanisms with its ability to positively augment signalling activation following ssRNA and CpG DNA detection in a host cell. Here we show that viperin interacts with the signalling adaptor proteins, STING and TBK1, which are central to the dsDNA signalling pathways, to enhance the activation of TBK1. In previous work viperin was shown to interact with the signalling adaptor protein IRAK1 which is central to the TLR7 & 9 signalling pathway [27]; However, in this instance viperin was also demonstrated to interact with the E3 ligase TRAF6 which was responsible for the polyubiquitination of IRAK1. While it is evident viperin directly binds to the adaptor protein STING (Figure 3E), which is the upstream adaptor protein of TBK1, the E3 ligase responsible for ligating the K63-linked ubiquitin chains to TBK1 remains unknown. Microscopy based analysis confirmed that viperin co-localises with the E3 ligase TRAF6 following dsDNA stimulation (Supplementary Figure 2A and B), in a similar manner to that seen following activation of TLR7 and TLR9 [21]. We were also able to show that this is the case for viperin and the E3 ligase TRAF3 following dsDNA stimulation (Supplementary Figure 2A and C). Both TRAF3 and TRAF6 have been previously implicated in the STING-TBK1 signalling axis [32, 33], however, further analysis is required to determine the functional relevance of this co-localisation to viperin’s enhancement of TBK1 activation.

The ability of viperin to enhance the dsDNA signaling pathway presents a novel mechanism for the protein’s antiviral capacity against DNA viruses. To investigate this in the absence of potential direct antiviral activity, we utilised a plasmid-based induction model of HBV replication in a hepatocyte cell line. In this HBV model system, overexpression of viperin resulted in enhanced activity of the IFN-β promoter in HepG2 cells transfected with both HBV genotypes D and A (Figure 5A), which correlated with a reduction in HBsAg and HBeAg present in the culture media (Figure 5B and 5C). Through viperin’s direct interaction with STING and enhancement of TBK1 activation, we postulate that viperin enhances the type-I interferon response to HBV DNA in this viral model, enabling enhanced cellular control of the viral infection *in vitro*. Together this data further highlights the importance of viperin’s interaction with adaptor molecules of the DNA signalling pathway to inhibit DNA viral infection.

Viperin’s ability to generate ddhCTP to inhibit the replication of members of the *Flavivirus* genus via inhibition of polymerase function, relies on its enzymatic cleavage of SAM [20]; however, ddhCTP was found to not inhibit the polymerases of members of the *Picornaviridae* family. Additionally, the ability of ddhCTP to inhibit the HCV RdRp was considerably lower than either of the *Flaviviruses*, DENV or WNV RdRps in an *ex vivo* assay [20], and other more diverse viruses such as HIV [10] and BUNV [49] are also inhibited by the functions of viperin’s enzymatic radical SAM domain in an as yet unspecified manner. Moreover, previous studies have demonstrated that the deletion of viperin’s N-terminus significantly abrogates its inhibition of not only HCV [14], but also CHIKV [50], and WNV [51]; a domain absent in the recombinant *Rattus norvegicus* viperin utilised to generate ddhCTP [20]. This evidence is suggestive of a potentially alternate antiviral role to viperin’s generation of ddhCTP but reliant on its radical SAM domain, to achieve the observed levels of viperin mediated inhibition of multiple viruses.

The radical SAM activity of viperin may contribute to its ability to enhance innate immune signalling and be regulated by the alternate [4Fe-4S] insertion mechanisms. Here we demonstrate viperin’s requirement for the insertion of the [4Fe-4S] cluster, a cofactor necessary for viperin’s enzymatic activity (Figure 1), within its radical SAM domain to enhance the type-I IFN response to dsDNA (Figure 6B). The insertion alone of this [4Fe-4S] cluster has previously been shown to stabilise viperin [52], and is primarily inserted by the cytosolic iron-sulphur protein assembly (CIA) targeting complex CIA1-CIA2B-MMS19, via binding to the C-terminal W361 residue of viperin (Figure 1)[24]. We have shown that the deletion of viperin’s C-terminus, and hence deletion of viperin’s interaction with this CIA complex, significantly increased viperin’s ability to enhance dsDNA signalling while mutation of viperin’s M1 domain which completely abrogates viperin’s [4Fe-4S] binding capacity conversely reduced viperin’s enhancement of dsDNA signalling (Figure 6B). This suggests that viperin’s [4Fe-4S] insertion is imperative to its ability to enhance dsDNA signalling, but that insertion of the [4Fe-4S] may not be solely the role of the CIA1-CIA2B-MMS19. Indeed the original findings describing viperin [4Fe-4S] insertion show a compensatory role of the alternate CIA targeting factor CIA2A to form a complex with CIA1 (CIA2A-CIA1) and contribute to viperin [4Fe-4S] insertion in cells lacking its alternate isoform CIA2B, albeit less efficiently than the CIA1-CIA2B-MMS19 complex [24]. As CIA2A binds to viperin’ N-terminus (Figure 1)[24], it is possible this protein maintains viperin’s [4Fe-4S] insertion in the C-terminally truncated viperin mutant which is unable to bind the CIA1-CIA2B-MMS19 complex (Figure 6B). However, the abrogation of viperin’s interaction with CIA2A through deletion of its N-terminus reduces its ability to enhance dsDNA signalling, suggests this interaction with CIA2A confers more than [4Fe-4S] insertion.

The less efficient contribution of CIA2A to viperin’s [4Fe-4S] insertion may lessen viperin’s enzymatic generation of 5’-dA to extend viperin’s enhancement of dsDNA signalling. Viperin generates 5’-dA as a by-product of its radical SAM enzymatic function through the utilisation of [4Fe-4S] clusters (Figure 1). High levels of 5’-dA have been demonstrated to act as a general inhibitor of radical SAM enzymes [48]. We have shown previously that when viperin and IRAK1/TRAF6 interact [47], high levels of 5’-dA are produced, and similarly, here we demonstrate for the first time, that this also occurs during viperin’s interaction with STING (Figure 7A). Concurrently, we also see viperin degradation when overexpressed with STING (Figure 7B), which is also the case when viperin is in the presence of overexpressed IRAK/TRAF6 [20, 47]. This would indicate the existence of a self-limiting negative feedback loop of viperin’s enhancement of innate immune signalling, whereby viperin’s enzymatic activity is promoted during its interaction with STING/IRAK1/TRAF6 until 5’-dA levels reach inhibitory concentrations and lead to the degradation of viperin protein. Moreover, as the CIA2A-CIA1 complex contributes to viperin [4Fe-4S] insertion less efficiently than the CIA1-CIA2B-MMS19 complex, presumably during CIA2A depletion, which reduces viperin’s ability to enhance the type-I IFN response following dsDNA recognition (Figure 8C), CIA2B would have no competition for binding to CIA1, and would efficiently facilitate generation of the self-limiting factor 5’-dA. Conversely, CIA2A overexpression leads to further enhancement of this pathway (Figure 8D). In this context the abundance of CIA2A likely outcompetes endogenous CIA2B for complex formation with CIA1. As CIA2A less efficiently contributes to the insertion of [4Fe-4S] into viperin and hence formation of 5’-dA, the overrepresented CIA2A likely mitigates viperin’s degradation and allows for prolonged enhancement of the dsDNA pathway. The dual engagement with these two distinct CIA isoforms appears to represent a novel regulatory mechanism of viperin’s antiviral activity, although further research is required to determine to contextual significance of each interaction.

The evolution of viperin predicates the protein’s role in innate immune regulation. Viperin is highly conserved, showing high amino acid identity across not only vertebrates including mammals, fish (reviewed in [7]) and reptiles [53], but also invertebrates such as oysters [54]. A recent study of the type-I ‘interferome’ identified viperin as a core IFN-induced antiviral factor across numerous vertebrate species [55], highlighting the protein’s ancestral role in antiviral innate immunity. Interestingly, a separate study of transcriptional divergence of the innate immune response between species revealed the high conservation of genes encoding proteins involved in immune response regulation as opposed to those with more direct acting effects on viral invasion [56]. Together this data provides evidence for viperin’s ancestral role as a regulator of the innate immune response to viral infection, and here we describe another instance of viperin’s enhancement of innate immune signalling events, complementary to its role in positively regulating TLR7/9 signalling (Figure 9)[27].

**Figure 9.**
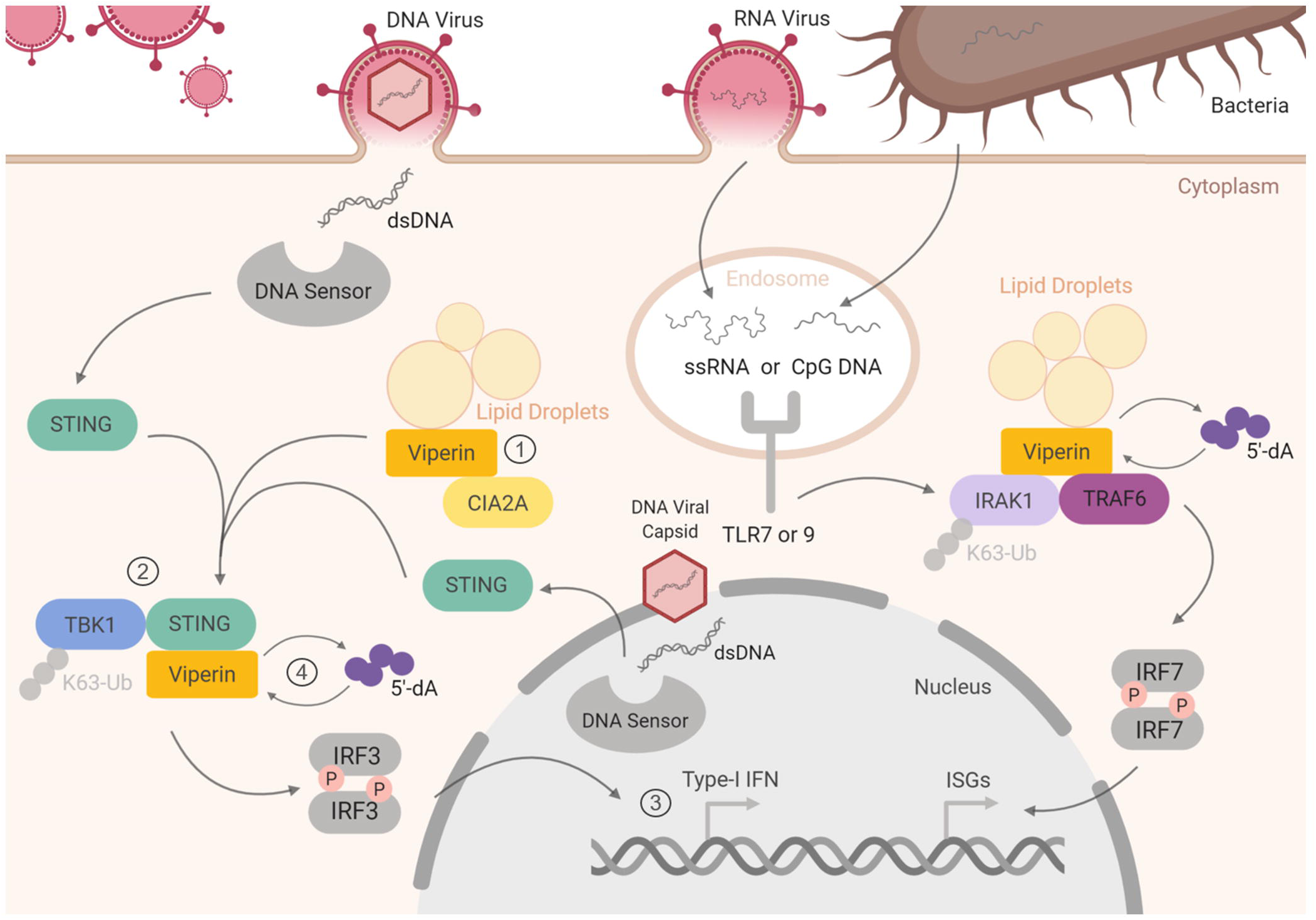
Viperin facilitates the formation of signalling enhancesomes to positively augment interferon production. Here we show that viperin can enhance the dsDNA signalling pathway (Image left) in a similar manner to its enhancement of the TLR7/9 signalling pathways (Image right). In summary we have shown that upon detection of aberrant dsDNA either within the cytosol or nucleus, viperin is able to enhance the dsDNA signalling pathway; 1) viperin pre-emptively associates with CIA2A, 2) viperin interacts with STING and enhances the activation of TBK1 through K63 polyubiquitination, 3) this process enhances the type-I interferon response to dsDNA and limits DNA viral infection, 4) viperin’s interaction with STING drives its enzymatic generation of 5’-dA, which results in viperin degradation.

The continued understanding of viperin’s antiviral activity offers valuable insight into the development of novel antiviral therapeutics. The study of viperin’s enzymatic function has already lead to the synthetic generation of the metabolic precursor of ddhCTP - ddhC – which shows promise as an antiviral therapeutic for its ability to enter human cells and inhibit the replication of ZIKV *in vitro* [20, 27]. A clearer understanding of viperin’s role in augmenting a multitude of diverse innate signalling events may additionally highlight complementary approaches to a broad range of small-molecule agonist therapies to combat antiviral therapeutics.

## Acknowledgements

This work was supported by the NHMRC (APP1053206) of Australia and a Research Focus Area small grant from La Trobe University. The authors would like to acknowledge the La Trobe University Microscopy Platform for their assistance in the microscopy aspects of this work; Associate Professor Anna Overby from Umea University for supplying CIA2A expression constructs, and Dr Michael Gantier for constructive comments.

## Author contributions

K.J. H., K.M.C and E.N.G.M. conceived the experiments; K.M.C., E.A.M., A.B.D., performed the experiments; M.S., K.H.V., P.A.R., E.N.G.M. and M.R.B. generated and/or supplied experimental reagents and materials; Y.T. and D.C.T. contributed to the interpretation of the results; K.M.C. and K.H. analysed the data and wrote the manuscript; all authors participated in reviewing and editing the final manuscript.

## Conflict of interest

The authors declare that they have no conflict of interest.

## Materials and Methods

### Cells and culture conditions

All mammalian cell lines were maintained at 37□°C in a 5% CO_2_ air atmosphere. Huh-7 human hepatoma cells, HeLa human epithelial cells, HEK293T human embryonic kidney cells, as well as primary murine embryonic fibroblast (MEF) cells were maintained in DMEM (Gibco) containing 10% (v/v) FCS. HepG2 human hepatoma cells were maintained in MEM (Gibco) containing 10% (v/v) FCS. The viperin^−/−^ MEFs were generated and prepared as previously described [9]. The polyclonal Huh-7 cell line stably expressing shRNA targeting viperin mRNA was as previously described [14].

### Real time PCR

All experiments involving real-time PCR were performed in 12-well plates with cells seeded at 7 × 10^4^ per well, 24 hrs prior to transfection. Total RNA was extracted from cells using TriSure reagent (Bioline), with first strand cDNA being synthesized from total RNA and reverse transcribed using a Tetro cDNA synthesis kit (Bioline). Quantitative real-time PCR was performed in a CFX Connect Real-Time Detection System (BioRad) to quantitate the relative levels of IFN and ISG mRNA in comparison to the house keeping gene RPLPO. Primers sequences were as follows: RPLOPO-FP 5’-AGA TGC AGC AGA TCC GCA T-3’, RPLPO-RP 5’-GGA TGG CCT TGC GCA-3’, IFI6-FP 5’-CCT GCT GCT CTT CAC TTG CA-3’, IFI6-RP 5’-CCG ACG GCC ATG AAG T-3’, OAS-FP 5’-TCC ACC TGC TTC ACA GAA CTA CA-3’, OAS-RP 5’-GGC GGA TGA GGC TCT TGA G-3’, mIFN-β-FP 5’-AGA AAG GAC GAA CAT TGG GAA A-3’, mIFN-β-RP 5’-TAG CAG AGC CCT TTT TGA TAA TGT AA-3’.

### Immunoprecipitation analysis

Where stated, prior to immunoprecipitation, cells were incubated with No-Weigh™ Format DSS crosslinker (Thermo-Fisher Scientific) for 30 mins at RT in ice-cold PBS (1.35mM DSS, pH 8.0), and then in quench solution (15 mM Tris, pH 7.5) for 15 mins at RT. Cell extracts were prepared with 0.5% (w/v) CHAPS lysis buffer supplemented with protease inhibition cocktail (Sigma). Lysates were pre-cleared with protein A/G PLUS-agarose beads (Santa Cruz Biotechnology) washed with 0.5% (w/v) CHAPS, immunoprecipitated with 2 µg/sample of indicated antibodies overnight at 4 °C before addition of washed protein A/G PLUS-agarose beads (Santa Cruz Biotechnology) for 1 hr with rotation at 4 °C. After extensive washes with the same lysis buffer, the immunoprecipitates were subject to immunoblot analysis.

### Immunoblot analysis

Lysates were subjected to SDS-PAGE. Proteins were transferred to 0.2 µm nitrocellulose membranes (Bio-Strategy) and probed with indicated primary antibodies. The protein bands were visualized using a SuperSignal West Femto Maximum Sensitivity Substrate (Thermo-Fisher Scientific) for horseradish peroxidase (HRP) conjugated secondary antibodies. The probing with the monoclonal mouse anti-β-actin antibody (Sigma) was used as a loading control. Membranes were scanned using an Amersham 600 chemiluminescent imager.

### Dual luciferase reporter assay

Luciferase experiments were performed essentially as previously described [57]. Cells were seeded at 4 × 10^4^ per well in 24-well plates, 24 hrs prior to transient transfection using Viafect (Promega) with 250 ng of a specified target construct as well as 250 ng pIFN-β-Firefly luciferase in combination with 2.5 ng of the constitutively expressing Renilla luciferase plasmid, pRL-TK. Following a further 24 hrs, cells were stimulated with synthetic viral mimics for specified time periods. Cells were lysed with 1 x PLB (Promega) and the luciferase outputs were measured with a dual luciferase reporter assay system (Promega) on a CLARIOstar (BMG LABTECH) microplate reader. All conditions were performed in at least triplicate.

### Viral infection and viral mimics

Transfection of HBV was performed as previously described [28], in HepG2 cells using a recombinant 1.3-mer transient transfection model system for HBV genotypes A and D. Cells were seeded at 1 × 10^5^ per well in 24-well plates, 24 hrs prior to transient transfection using Viafect (Promega) with a combined total of 500 ng per well of the specified HBV 1.3 mer plasmid as well as target and luciferase plasmids. Cell supernatants were harvested at specified time points for quantitative serology as previously described [42], and cell lysates were harvested for dual luciferase reporter assays.

Herpes simplex virus type I (HSV-1) strain KOS was used to infect HeLa cells washed with phosphate-buffered saline (PBS) at the indicated multiplicity of infection (MOI) in DMEM without FCS. After 1 hr of incubation, the infection medium was removed and replaced with DMEM containing 2.5% (v/v) FCS.

The dsDNA viral mimic poly dA:dT (Invivogen) was transfected into cells using DMRIE-C reagent (Life Technologies) as per manufacturer’s instructions at a concentration of 1□µg/ml.

### Immunofluorescence microscopy

All immunofluorescence staining was performed as previously described [58], and was visualised using either a Nikon Eclipse T*i*-E fluorescence inverted microscope or a Ziess Confocal LSM 780 microscope, and images were captured using NIS Elements software or ZEN microscopy software respectively.

### Proximity ligation assay

Cells were seeded at 7 × 10^4^ per well in 12-well plates, 24 hrs prior to transient transfection using Viafect (Promega) with the specified viperin-flag constructs. Following a further 24 hrs, cells were trypsinised and seeded at 3 × 10^3^ per well in a 96-well plate, allowed to recover and then stimulated with viral mimics. Cells were fixed with 4% (w/v) paraformaldehyde and the proximity ligation assay (PLA) was conducted using the Duolink® In Situ Kit (Merck) as per the manufacturer’s instructions. Positive interactions were visualized using a Nikon Eclipse T*i*-E fluorescence inverted microscope and images were captured using NIS Elements software.

### Enzyme activity assay

HEK293T cells transfected with viperin, and/or STING and TBK1 were harvested from one 10 cm diameter tissue culture plate each, resuspended in 500 µl of anoxic Tris-buffered saline (50 mM Tris-Cl, pH 7.6, 150 mM NaCl) containing 1% Triton X-100, sonicated within an anaerobic glovebox (Coy Chamber), and centrifuged at 14,000 g for 10 min. Dithiothreitol (DTT; 5 mM) and dithionite (5 mM) were added to the cell lysate together with CTP (300 μM). The assay mixture was incubated at room temperature for 30 min prior to starting the reaction by the addition of SAM (200 µM). The assay was incubated for 60 min at room temperature, after which the reaction was stopped by heating at 95 °C for 10 min. The solution was chilled to 4 °C, and the precipitated proteins were removed by centrifugation at 14,000 rpm for 25 min. The supernatant was then extracted with acetonitrile. The amount of 5’-dA in the samples was quantified by UPLC-tandem mass spectrometry, and the amount of viperin present in the lysate quantified by immunoblotting as described previously [47]. Results reported represent the average of 3 biological replicates of the assay.

### Generation of CIA2A knockout HeLa cell lines

The *CIA2A*-targeting lentivirus was packaged using lentiCRISPRv2 system. HeLas were seeded at 2 × 10^5^ per well in a 6-well plate and were transduced the next day with the lentiviral particles expressing *Cas9* and three independent *CIA2A*-specific guide RNAs (#1 5’-CAGCGTCCAGGAGAGCAGCC-3’, #2 5’-GGACGCTGAGCAGAGTCCTG-3’ and #3 5’-GGGCAGCTCCCGGCTCAGAG-3’). At 48 hr-post transduction, fresh media containing 0.5 μg/ml puromycin was added to select for cells transduced with the *CIA2A*-targeting lentivirus and *Cas9* expression. After culturing in puromycin-containing media for an additional 72 hours, the cells were expanded, and then analysed for CIA2A protein expression by immunoblot analysis with rabbit polyclonal anti-FAM96A(CIA2A) (Thermo-Fisher Scientific, REF#PA5-66628) antibody.

### Plasmid constructs and transfections

All plasmid constructs were transiently transfected into the indicated cells using Viafect Transfection Reagent (Promgea) as per manufacturer’s instructions. Viperin-GFP is within the pEGFP-C1 (Clontech) backbone where GFP is expressed as an N-terminal fusion protein. The corresponding control was empty pEGFP-C1. Viperin-flag consists of the pFLAG-CMV™ Gateway® backbone (Invitrogen), where a CMV promoter drives expression of the viperin with a flag tag at the N-terminus. The viperin mutant plasmids viperin-5’Δ33-flag, viperin-3’Δ17-flag and Viperin-SAM1-flag, were created as previously described [14]. Viperin-mCherry and mCherry plasmids were created as previously described [13]. The reporter plasmid IFN-β-Luc (IFN-Beta_pGL3) was a gift from Nicolas Manel (Addgene plasmid # 102597) [59]. pRL-TK (Promega) consists of a TK promoter which drives the constitutive expression of *Renilla* luciferase. The, pCMV-STING-3Xmyc and pCMV-TBK1-myc plasmids were a gift from Russell Diefenbach (Westmead Millennium Institute, Sydney). These are vectors expressing an N-terminally myc-tagged STING or TBK1 proteins in transfected mammalian cells and are expressed constitutively by a CMV promoter. TBK1-mCherry was within pLenti-V5-D-TOPO-mCherry backbone where a CMV promoter drives its constitutive expression. The NS5A-TN50-viperin (NS5A-viperin) plasmid, which replaces the amphipathic helix of viperin with that of HCV NS5A was created as previously described [46]. The pEF-CIA2A-myc-his plasmid was a kind gift from Anna Överby (Umeå University, Sweden), and expresses N-terminally myc/His tagged CIA2A under the control of the EF-1α promoter. The ubiquitin plasmids pRK5-Ubiqutin-K48-HA and pRK5-Ubiquitin-K63-HA (Addgene) are constructed with a pRK5-HA backbone where the HA tag is at the N-terminus of ubiquitin. Hepatitis B virus 1.3 mer plasmid constructs for both genotype A and D are as previously described [42].

The Nucleofector transfection kit (Lonza) was used to transfect siRNA into cells, according to the manufacturer’s guide. In brief, 1 × 10^6^ of HeLa cells were transfected with 50 nM of siRNA universal control (Sigma Aldrich) or siRNA specific for viperin (SASI_Hs02_00362416; Sigma Aldrich) by a Nucleofector 2b Device (Lonza). Post 24 hr transfection, cells were used for infection or collected for immunoblot analysis. The sequence of the siRNA specific for viperin was 5’-AGA GCG GAA AGT GGA ACG AGA-3’.

### Statistical analysis

Results are expressed as mean ± S.E.M. Student’s t test was used for statistical analysis, with p < 0.05 considered to be significant. All statistical analysis was performed using Prism 8 (GraphPad Software).

## Figure legends

**Supplementary Figure 1.**

**(A)** Huh-7 cells were transfected with viperin-mCherry or control-mCherry and TBK1-myc constructs 24 hrs prior to stimulation with poly dA:dT for 2 hrs. Cell extracts were immunoprecipitated with rabbit monoclonal anti-mCherry antibody (Biovision) and subject to immunoblot analysis with indicated antibodies. **(B)** Huh-7 cells were transfected with control-mCherry, viperin-wt-mCherry, viperin-5’Δ33-mCherry or viperin-3’Δ17-mCherry and STING-myc constructs 24 hrs. Cell extracts were immunoprecipitated with rabbit monoclonal anti-mCherry antibody (Biovision) and subject to immunoblot analysis with indicated antibodies. Immunoblots are representative of at least 2 independent experiments.

**Supplementary Figure 2.**

**(A)** Huh-7 and were transfected with a viperin-flag construct 24 hrs prior to poly dA:dT stimulation for 2hrs. Cells were probed with mouse monoclonal anti-flag (Sigma) and rabbit monoclonal anti-TRAF6 (Cell Signalling) or anti-TRAF3 (Cell Signalling) antibodies, then subject to Duolink® *In Situ* Red Mouse/Rabbit PLA and DAPI staining. Imaged on Nikon Eclipse T*i*-E fluorescence inverted microscope. Scale bar represents 100 µm. Original magnification is X20. **(B & C)** HeLa cells were transfected with viperin-flag construct 24 hrs prior to poly dA:dT stimulation for 2hrs. Cells were subject to immunofluorescence staining with mouse monoclonal anti-flag (Sigma) and **(B)** rabbit monoclonal anti-TRAF6 (Cell Signalling) or **(C)** anti-TRAF3 (Cell Signalling) antibodies followed by an Alexa555-conjugated goat anti-mouse (Invitrogen) and Alexa488-conjugated goat anti-rabbit (Invitrogen) secondary, as well as DAPI staining. Imaged on Nikon Eclipse T*i*-E fluorescence inverted microscope. Scale bar represents 15 µm. Original magnification is X60.

## Notes

#### Summary of Updates

Additional data figures have been added, which (1) describes the radical SAM enzymatic activity of viperin in the context of its ability to interact with the signalling adapter molecules of the dsDNA signalling pathway (Figure 7), and (2) describes viperin’s interaction with the cytosolic iron-sulphur assembly protein CIA2A in the context of its ability to enhance dsDNA signalling (Figure 8). Consequently, multiple hypotheses from our original discussion have now been substantiated, and as such, the discussion in this revised submission has been extensively altered to include the new data sets. Additionally, Figures 1 and 9 have been added as schematic representations to aid the readers understanding of the research content covered in this revised submission.

## References

1. O’Neill LA, Bowie AG (2010) Sensing and signalling in antiviral innate immunity. Curr Biol.

2. Akira S, Uematsu S, Takeuchi O (2006) Pathogen recognition and innate immunity. Cell.

3. Goubau D, Deddouche S, Reis e Sousa C (2013) Cytosolic Sensing of Viruses. Immunity.

4. Stark GR, Darnell JE (2012) The JAK-STAT Pathway at Twenty. Immunity.

5. Schneider WM, Chevillotte MD, Rice CM (2014) Interferon-Stimulated Genes: A Complex Web of Host Defenses. Annu Rev Immunol 32: 513–545.

6. Helbig KJ, Beard MR (2014) The role of viperin in the innate antiviral response. J Mol Biol.

7. Helbig KJ, Beard MR (2014) The role of viperin in the innate antiviral response. J Mol Biol 426: 1210–1219.

8. Chin KC, Cresswell P (2001) Viperin (cig5), an IFN-inducible antiviral protein directly induced by human cytomegalovirus. Proc Natl Acad Sci U S A.

9. Van Der Hoek KH, Eyre NS, Shue B, Khantisitthiporn O, Glab-Ampi K, Carr JM, Gartner MJ, Jolly LA, Thomas PQ, Adikusuma F, et al. (2017) Viperin is an important host restriction factor in control of Zika virus infection. Sci Rep.

10. Nasr N, Maddocks S, Turville SG, Harman AN, Woolger N, Helbig KJ, Wilkinson J, Bye CR, Wright TK, Rambukwelle D, et al. (2012) HIV-1 infection of human macrophages directly induces viperin which inhibits viral production. Blood.

11. Wang X, Hinson ER, Cresswell P (2007) The Interferon-Inducible Protein Viperin Inhibits Influenza Virus Release by Perturbing Lipid Rafts. Cell Host Microbe.

12. Tang HB, Lu ZL, Wei XK, Zhong TZ, Zhong YZ, Ouyang LX, Luo Y, Xing XW, Liao F, Peng KK, et al. (2016) Viperin inhibits rabies virus replication via reduced cholesterol and sphingomyelin and is regulated upstream by TLR4. Sci Rep.

13. Helbig KJ, Carr JM, Calvert JK, Wati S, Clarke JN, Eyre NS, Narayana SK, Fiches GN, McCartney EM, Beard MR (2013) Viperin Is Induced following Dengue Virus Type-2 (DENV-2) Infection and Has Anti-viral Actions Requiring the C-terminal End of Viperin. PLoS Negl Trop Dis.

14. Helbig KJ, Eyre NS, Yip E, Narayana S, Li K, Fiches G, Mccartney EM, Jangra RK, Lemon SM, Beard MR (2011) The antiviral protein viperin inhibits hepatitis C virus replication via interaction with nonstructural protein 5A. Hepatology.

15. Seo JY, Yaneva R, Cresswell P (2011) Viperin: A multifunctional, interferon-inducible protein that regulates virus replication. Cell Host Microbe.

16. Broderick WE, Broderick JB (2019) Radical SAM enzymes: surprises along the path to understanding mechanism. J Biol Inorg Chem.

17. Holliday GL, Akiva E, Meng EC, Brown SD, Calhoun S, Pieper U, Sali A, Booker SJ, Babbitt PC (2018) Atlas of the Radical SAM Superfamily: Divergent Evolution of Function Using a “Plug and Play” Domain. In, Methods in Enzymology.

18. Landgraf BJ, McCarthy EL, Booker SJ (2016) Radical S-Adenosylmethionine Enzymes in Human Health and Disease. Annu Rev Biochem.

19. Wang J, Woldring RP, Román-Meléndez GD, McClain AM, Alzua BR, Marsh ENG (2014) Recent advances in radical SAM enzymology: New structures and mechanisms. ACS Chem Biol.

20. Gizzi AS, Grove TL, Arnold JJ, Jose J, Jangra RK, Garforth SJ, Du Q, Cahill SM, Dulyaninova NG, Love JD, et al. (2018) A naturally occurring antiviral ribonucleotide encoded by the human genome. Nature.

21. Shaveta G, Shi J, Chow VTK, Song J (2010) Structural characterization reveals that viperin is a radical S-adenosyl-l-methionine (SAM) enzyme. Biochem Biophys Res Commun.

22. Duschene KS, Broderick JB (2010) The antiviral protein viperin is a radical SAM enzyme. FEBS Lett.

23. Stehling O, Mascarenhas J, Vashisht AA, Sheftel AD, Niggemeyer B, Rösser R, Pierik AJ, Wohlschlegel JA, Lill R (2013) Human CIA2A-FAM96A and CIA2B-FAM96B integrate iron homeostasis and maturation of different subsets of cytosolic-nuclear iron-sulfur proteins. Cell Metab.

24. Upadhyay AS, Stehling O, Panayiotou C, Rösser R, Lill R, Överby AK (2017) Cellular requirements for iron-sulfur cluster insertion into the antiviral radical SAM protein viperin. J Biol Chem.

25. Proud D, Turner RB, Winther B, Wiehler S, Tiesman JP, Reichling TD, Juhlin KD, Fulmer AW, Ho BY, Walanski AA, et al. (2008) Gene expression profiles during in vivo human rhinovirus infection insights into the host response. Am J Respir Crit Care Med.

26. Crosse KM, Monson EA, Beard MR, Helbig KJ (2018) Interferon-Stimulated Genes as Enhancers of Antiviral Innate Immune Signaling. J Innate Immun 10:.

27. Saitoh T, Satoh T, Yamamoto N, Uematsu S, Takeuchi O, Kawai T, Akira S (2011) Antiviral protein viperin promotes toll-like receptor 7- and toll-like receptor 9-mediated type i interferon production in plasmacytoid dendritic cells. Immunity.

28. Diebold SS, Kaisho T, Hemmi H, Akira S, Reis E Sousa C (2004) Innate Antiviral Responses by Means of TLR7-Mediated Recognition of Single-Stranded RNA. Science (80-).

29. Lund JM, Alexopoulou L, Sato A, Karow M, Adams NC, Gale NW, Iwasaki A, Flavell RA (2004) Recognition of single-stranded RNA viruses by Toll-like receptor 7. Proc Natl Acad Sci U S A.

30. Ishikawa H, Barber GN (2008) STING is an endoplasmic reticulum adaptor that facilitates innate immune signalling. Nature.

31. Ishikawa H, Ma Z, Barber GN (2009) STING regulates intracellular DNA-mediated, type i interferon-dependent innate immunity. Nature.

32. Zhong B, Yang Y, Li S, Wang YY, Li Y, Diao F, Lei C, He X, Zhang L, Tien P, et al. (2008) The Adaptor Protein MITA Links Virus-Sensing Receptors to IRF3 Transcription Factor Activation. Immunity.

33. Saitoh T, Fujita N, Hayashi T, Takahara K, Satoh T, Lee H, Matsunaga K, Kageyama S, Omori H, Noda T, et al. (2009) Atg9a controls dsDNA-driven dynamic translocation of STING and the innate immune response. Proc Natl Acad Sci U S A.

34. Tanaka Y, Chen ZJ (2012) STING specifies IRF3 phosphorylation by TBK1 in the cytosolic DNA signaling pathway. Sci Signal.

35. Mukai K, Konno H, Akiba T, Uemura T, Waguri S, Kobayashi T, Barber GN, Arai H, Taguchi T (2016) Activation of STING requires palmitoylation at the Golgi. Nat Commun.

36. Heaton SM, Borg NA, Dixit VM (2016) Ubiquitin in the activation and attenuation of innate antiviral immunity. J Exp Med.

37. Ni G, Konno H, Barber GN (2017) Ubiquitination of STING at lysine 224 controls IRF3 activation. Sci Immunol.

38. Wang Q, Liu X, Cui Y, Tang Y, Chen W, Li S, Yu H, Pan Y, Wang C (2014) The E3 Ubiquitin ligase AMFR and INSIG1 bridge the activation of TBK1 kinase by modifying the adaptor STING. Immunity.

39. Liu S, Cai X, Wu J, Cong Q, Chen X, Li T, Du F, Ren J, Wu YT, Grishin N V., et al. (2015) Phosphorylation of innate immune adaptor proteins MAVS, STING, and TRIF induces IRF3 activation. Science (80-).

40. Ouyang S, Song X, Wang Y, Ru H, Shaw N, Jiang Y, Niu F, Zhu Y, Qiu W, Parvatiyar K, et al. (2012) Structural Analysis of the STING Adaptor Protein Reveals a Hydrophobic Dimer Interface and Mode of Cyclic di-GMP Binding. Immunity.

41. Tu D, Zhu Z, Zhou AY, Yun C hong, Lee KE, Toms A V., Li Y, Dunn GP, Chan E, Thai T, et al. (2013) Structure and Ubiquitination-Dependent Activation of TANK-Binding Kinase 1. Cell Rep.

42. Sozzi V, Walsh R, Littlejohn M, Colledge D, Jackson K, Warner N, Yuen L, Locarnini SA, Revill PA (2016) In Vitro Studies Show that Sequence Variability Contributes to Marked Variation in Hepatitis B Virus Replication, Protein Expression, and Function Observed across Genotypes. J Virol.

43. Guo F, Tang L, Shu S, Sehgal M, Sheraz M, Liu B, Zhao Q, Cheng J, Zhao X, Zhou T, et al. (2017) Activation of stimulator of interferon genes in hepatocytes suppresses the replication of hepatitis B virus. Antimicrob Agents Chemother 61:.

44. He J, Hao R, Liu D, Liu X, Wu S, Guo S, Wang Y, Tien P, Guo D (2016) Inhibition of hepatitis B virus replication by activation of the cGAS-STING pathway. J Gen Virol 97: 3368–3378.

45. Upadhyay AS, Vonderstein K, Pichlmair A, Stehling O, Bennett KL, Dobler G, Guo JT, Superti-Furga G, Lill R, Överby AK, et al. (2014) Viperin is an iron-sulfur protein that inhibits genome synthesis of tick-borne encephalitis virus via radical SAM domain activity. Cell Microbiol.

46. Makins C, Ghosh S, Román-Meléndez GD, Malec PA, Kennedy RT, Marsh ENG (2016) Does viperin function as a radical S-adenosyl-L-methioninedependent enzyme in regulating farnesylpyrophosphate synthase expression and activity? J Biol Chem.

47. Dumbrepatil AB, Ghosh S, Zegalia KA, Malec PA, Hoff JD, Kennedy RT, Marsh ENG (2019) Viperin interacts with the kinase IRAK1 and the E3 ubiquitin ligase TRAF6, coupling innate immune signaling to antiviral ribonucleotide synthesis. J Biol Chem.

48. Choi-Rhee E, Cronan JE (2005) A nucleosidase required for in vivo function of the S-adenosyl-L-methionine radical enzyme, biotin synthase. Chem Biol.

49. Carlton-Smith C, Elliott RM (2012) Viperin, MTAP44, and Protein Kinase R Contribute to the Interferon-Induced Inhibition of Bunyamwera Orthobunyavirus Replication. J Virol.

50. Teng TS, Foo SS, Simamarta D, Lum FM, Teo TH, Lulla A, Yeo NKW, Koh EGL, Chow A, Leo YS, et al. (2012) Viperin restricts chikungunya virus replication and pathology. J Clin Invest.

51. Jiang D, Weidner JM, Qing M, Pan X-B, Guo H, Xu C, Zhang X, Birk A, Chang J, Shi P-Y, et al. (2010) Identification of Five Interferon-Induced Cellular Proteins That Inhibit West Nile Virus and Dengue Virus Infections. J Virol.

52. Haldar S, Paul S, Joshi N, Dasgupta A, Chattopadhyay K (2012) The presence of the iron-sulfur motif is important for the conformational stability of the antiviral protein, viperin. PLoS One.

53. Milic NL, Davis S, Carr JM, Isberg S, Beard MR, Helbig KJ (2015) Sequence analysis and characterisation of virally induced viperin in the saltwater crocodile (Crocodylus porosus). Dev Comp Immunol.

54. Green TJ, Speck P, Geng L, Raftos D, Beard MR, Helbig KJ (2015) Oyster viperin retains direct antiviral activity and its transcription occurs via a signalling pathway involving a heat-stable haemolymph protein. J Gen Virol.

55. Shaw AE, Hughes J, Gu Q, Behdenna A, Singer JB, Dennis T, Orton RJ, Varela M, Gifford RJ, Wilson SJ, et al. (2017) Fundamental properties of the mammalian innate immune system revealed by multispecies comparison of type I interferon responses. PLoS Biol.

56. Hagai T, Chen X, Miragaia RJ, Rostom R, Gomes T, Kunowska N, Henriksson J, Park JE, Proserpio V, Donati G, et al. (2018) Gene expression variability across cells and species shapes innate immunity. Nature.

57. Helbig KJ, George J, Beard MR (2005) A novel I-TAC promoter polymorphic variant is functional in the presence of replicating HCV in vitro. J Clin Virol.

58. Narayana SK, Helbig KJ, McCartney EM, Eyre NS, Bull RA, Eltahla A, Lloyd AR, Beard MR (2015) The interferon-induced transmembrane proteins, IFITM1, IFITM2, and IFITM3 inhibit hepatitis C virus entry. J Biol Chem.

59. Gentili M, Kowal J, Tkach M, Satoh T, Lahaye X, Conrad C, Boyron M, Lombard B, Durand S, Kroemer G, et al. (2015) Transmission of innate immune signaling by packaging of cGAMP in viral particles. Science (80-).

