## Supplementary figures and images for "Viperin binds STING and enhances the type-I interferon response following dsDNA detection"

### Supplementary Figure 1

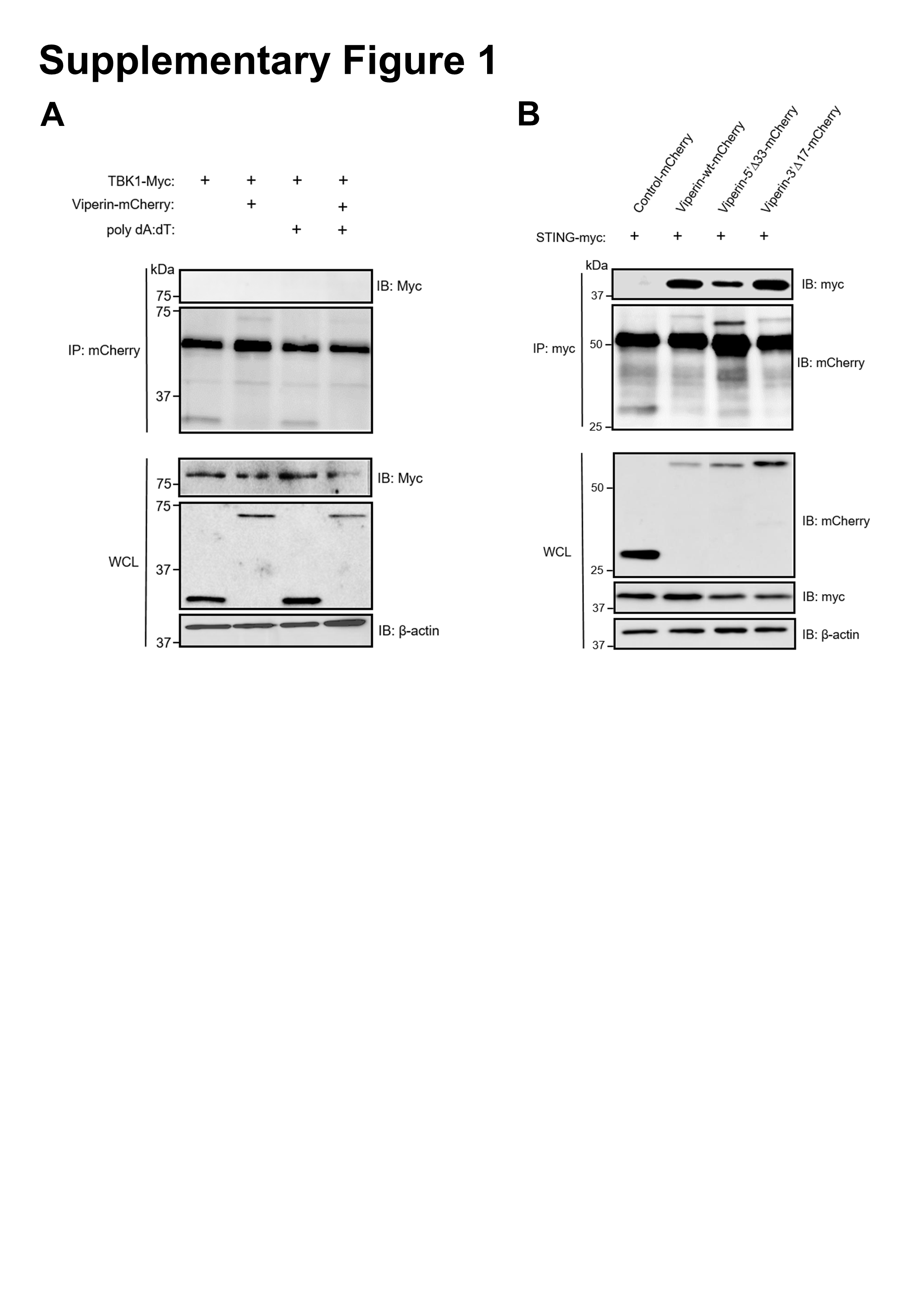

### Supplementary Figure 2

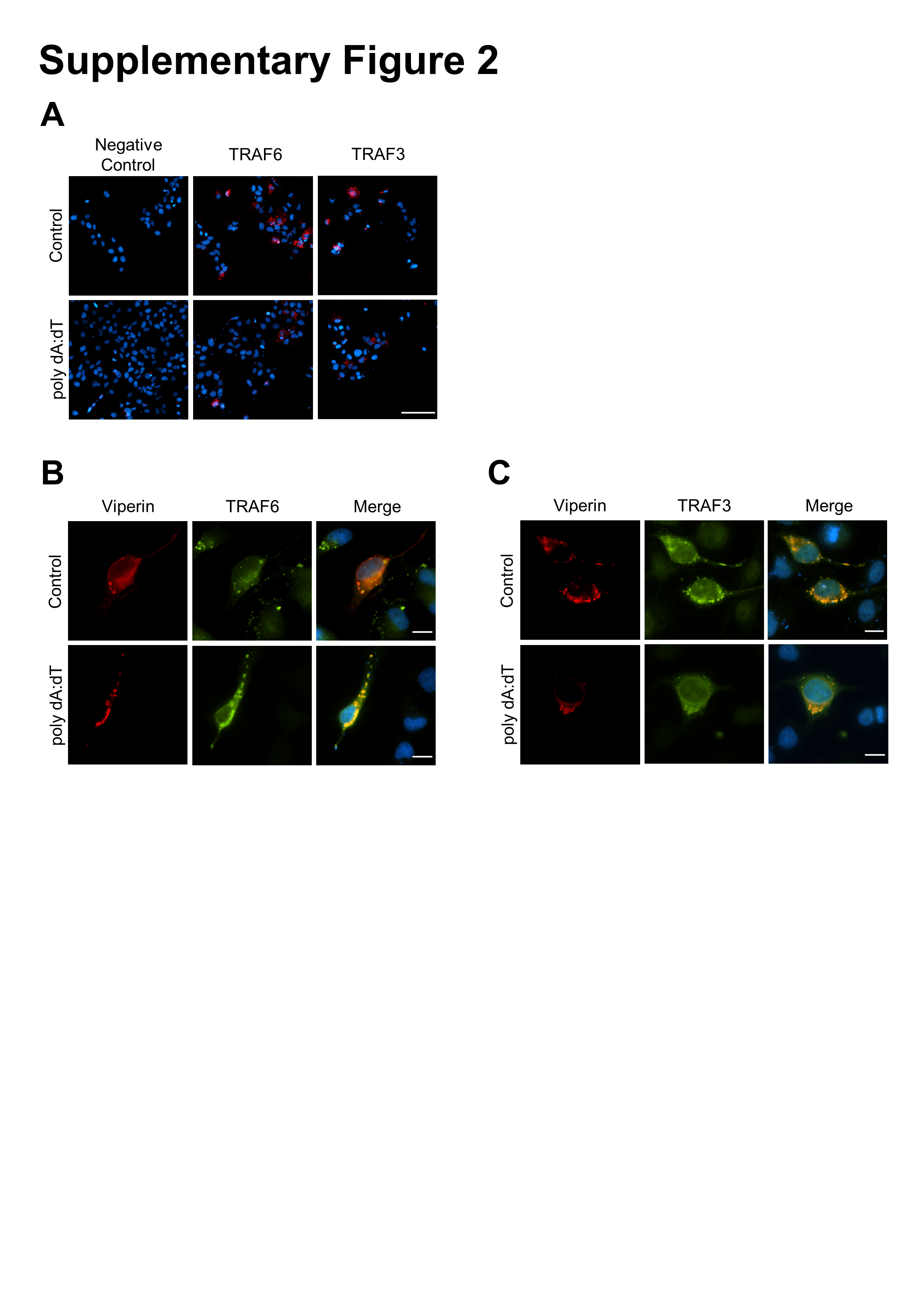
